# Topical Delivery of 4-Aminopyridine Enhances Skin Regeneration in Burn Wounds

**DOI:** 10.1101/2025.06.05.658142

**Authors:** Govindaraj Ellur, Meenakshi Kamaraj, John C. Elfar, Johnson V John, Prem Kumar Govindappa

## Abstract

Burn wounds are a common traumatic injury that impair cellular function and hinder the healing process, often resulting in significant skin loss. While autologous skin grafting is considered the gold standard for treating burns, its widespread use is limited due to donor site morbidity and the requirement for large amounts of tissue. Traditional wound dressings and treatments often fail to ensure complete recovery. Being initially FDA-approved to treat multiple sclerosis, 4-aminopyridine (4-AP) has also been shown to accelerate burn wound closure by transforming keratinocytes and fibroblasts when administered systemically. However, prolonged systemic use of 4-AP can lead to significant side effects. In this study, we aimed to repurpose 4-AP for treating skin burn wounds by delivering it topically using a laponite-gelatin gel formulation. This method allows for non-invasive and localized drug delivery on burn wound site. We evaluated the physical properties of the 4-AP gel shear thinning behavior, drug release kinetics, biocompatibility, and functional wound closure using a scratch assay. Moreover, our *in vivo* experiments showed that the 4-AP loaded gel accelerates wound healing by enhancing re-epithelialization and hair follicle regeneration and promoting fibroblast to myofibroblast transformation, which supports extracellular matrix remodeling after skin burns. This novel application of the 4-AP gel could offer a promising alternative to current burn wound therapies, potentially leading to improved outcomes for burn patients.

## Introduction

Wounds such as diabetic ulcers, burn injuries, and pressure ulcers show unique healing challenges due to their distinct etiologies and healing environments [1–3]. Among them, burn injuries are particularly critical, due to its extensive tissue damage, high risk of infection, prolonged inflammation, fluid loss, and impaired vascularization, all of which complicate tissue regeneration and increase morbidity and mortality [4–7]. Specifically, burn wounds represent a prevalent and challenging burden in critical care, often requiring complex, long-term management strategies [2]. In the U.S., acute thermal injuries affect close to 500,000 people annually, resulting in around 40,000 hospitalizations and 3,400 deaths [8]. Over the past few decades, significant advancements have been made in developing novel approaches to treat burn injuries with different forms of materials including the non-adherent films, gauzes, foams, alginates, hydrocolloids, and hydrogels, each with unique properties to address different burn wound needs [9–12]. Severe thermal skin burns are often exacerbated by prolonged inflammation and increased cell death, leading to delayed wound healing and substantial morbidity. Interestingly, our previous study on repurposing the systemic administration of 4-aminopyridine (4-AP), an FDA-approved non-specific potassium channel blocker for multiple sclerosis [13, 14], showed its therapeutic potential in a murine model of severe burn injury. Early administration of 4-AP significantly reduced pro-inflammatory cytokines IL-1β and TNF-α, while concurrently increasing anti-inflammatory markers such as CD206, ARG-1, and IL-10, suggesting a favorable shift in the inflammatory response during the initial phase of wound healing. The systemic administration of 4-AP accelerates healing in full-thickness excisional and burn wounds [15, 16]. The promising results from these studies led us to FDA exemptions for conducting clinical trials on excisional and burn wounds (ClinicalTrials.gov IDs: NCT06333171 and NCT06596434).

While systemic administration of 4-AP has shown efficacy, however there is growing interest in developing a minimally invasive localized delivery methods to minimize potential side effects and optimize therapeutic outcomes [17]. Topical application of drugs offers several advantages, including minimally invasive delivery, reduced systemic exposure, and potentially improved patient compliance [18, 19]. However, the development of an effective topical formulation requires careful consideration of factors such as drug release kinetics, stability of the gel, biocompatibility, and physical properties of the delivery vehicle [20, 21]. In this context, injectable hydrogels have emerged as versatile platforms for topical drug delivery [22, 23]. Their high-water content, shear-thinning characteristic, biocompatibility, and tunable physical properties make them ideal candidates for wound healing applications [24]. However, their long-term stability and rapid degradation rates can limit clinical utility. Recently, silica nanoplatelets commonly referred to as Laponite® have been investigated for their enhanced shear-thinning properties in hydrogel compositions [25]. Recent studies on Laponite/gelatin composed gel formulations have demonstrated sustained drug release and favorable injectability, including shear-thinning behavior upon administration through needles or catheters for various biomedical applications [26–28]. Later these compositions became a commercially available product such as Obsidio™ (Boston Scientific), an injectable soft solid for embolization derived from Laponite. Specifically, the use of natural and synthetic components such as composite hydrogels have shown promise in achieving controlled drug release and promoting tissue regeneration [29–31].

In this study we are re-purposing the two FDA approved products for burn wound healing studies and formulated a composite hydrogel by combining Laponite and gelatin. This combinatorial physical crosslinking strategy significantly influences the 4-AP release kinetics and imparts the shear-thinning characteristics necessary for effective topical gel application. The 4-AP loaded 4-AP gel-administered at lower doses every three days could enhance burn wound healing compared to systemic high-dose daily treatment reported from us previously [16]. To achieve this goal, we first focused on optimizing various gel formulations by adjusting gelatin and laponite concentrations. Thus, we conducted comprehensive rheological analyses to characterize the physical properties of these formulations, with particular attention to their shear-thinning behavior and mechanical stability. These properties are crucial for ensuring ease of application and maintaining structural integrity at the wound site. Following the optimization of the gel formulation, we evaluated its drug release profile and biocompatibility. *In vitro* wound model studies were conducted to assess the 4-AP gel’s potential to promote cell migration and wound closure. Finally, we investigated the effect of the 4-AP gel *in vivo* using a burn wound model, examining its benefits on wound closure, tissue regeneration, and key molecular markers associated with wound healing. By exploring this localized, low-dose treatment strategy, we seek to offer a less invasive alternative to systemic treatments, potentially improving patient outcomes and expanding the therapeutic options available for managing challenging wound conditions.

## Materials and Methods

### 4-aminopyridine loaded gel formulation and characterization

Laponite-Gelatin (Gel Control) and 4-aminopyridine (4-AP) drug-loaded laponite-gelatin (4-AP Gel) gels were formulated with slight modifications from existing developed protocols [26, 27]. Initially, stock gels of 10% w/v laponite (Laponite XLG, BYK) and 20% w/v gelatin (#G9382, Gelatin derived from porcine skin, Sigma) were prepared using Milli Q deionized (DI) water. Varied concentrations of laponite and gelatin gels were formulated by mixing them at specific ration mentioned in **Table 1**. The mixing was carried out at 3000 rpm using a DAC 150.1 FV-K, FlackTek, speed mixer for a duration of 5 min then incubation on ice for 5 min for 3 consecutive cycles. 4-AP concentration was kept at 20 μg per 1g of gel in case of drug-loaded gel (4-AP), which had DI water dissolved 4-APadded to the gel during the final mixing cycle to achieve homogenous distribution. The prepared gels were stored at 4 °C for 24 h before use and equilibrated to room temperature before any experimental procedures.

**Table 1.**
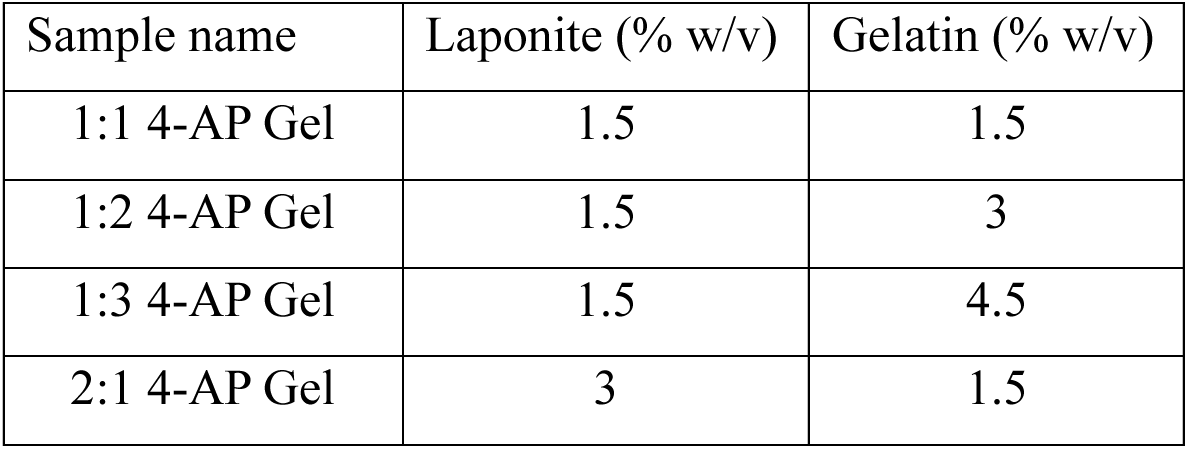
4-AP-loaded to varied concentrations of laponite and gelatin gels.

As prepared samples were taken for Fourier transform infrared spectroscopy (FTIR, Alpha II, Bruker). For FTIR analysis, wet gel samples were lyophilized and employed for peak identification and spectra was documented at 4 cm^−1^ spectral resolution over the 4000–800 cm^−1^ range. These developed gel formulations were employed for drug release study to understand their release kinetics. 500μg of gel was placed in transwell inserts and placed inside a 24-well plate and 1X DPBS was utilized as a release medium. Drug released into the 1X DPBS was quantified by measuring absorbance at 260nm for 4-AP using spectrophotometer (Biophotometer plus, Effendorf).

### Rheological properties analysis

The rheological behavior of the formulated gels was evaluated using a rheometer (MCR series, Anton Paar) equipped with an 8 mm parallel plate setup at 25 °C, following previously established protocols [32]. Strain-dependent storage moduli were determined using amplitude sweep measurements, with strain ranging from 1% to 100% at a constant frequency of 1 Hz. The viscosity of the formulated gels was measured across shear rates from 0.1 to 10 s⁻¹ to better assess the degree of injectability. For frequency sweep analysis, a constant strain of 1% was applied while varying the frequency from 0.01 to 100 Hz

### *In vitro* cell culture and biocompatibility assays

Human Dermal Fibroblasts (HDFs, PCS-201-012) were cultured in Dulbecco’s Modified Eagle Medium + GlutaMAX™ (#10-569-044, DMEM, Gibco) with 10 % fetal bovine serum and 1 % penicillin streptomycin (#15-140-163, Gibco) at 37 °C with 5 % CO_2_ humidified atmosphere. Once the cells reached the confluency, they were trypsinized using TrypLE™ (#12-605-036, Sigma) and utilized for future experiments. Similarly, Human Keratinocytes (HaCaTs) were cultured in DMEM complete media and trypsinized using 0.1% Trypsin-EDTA (#25300062, Sigma) and seeded for further experiments. Both cell types such as HDFs and HaCaTs were utilized for *in vitro* cell culture experiments to determine biomaterial cytotoxicity. For biocompatibility studies, cells were seeded at a density of 20000 cells per well in a 24-well plate and kept them at 37 °C with 5 % CO_2_ incubator for 1 h for cell to attach before exposing them to formulated gels. Following cell attachment, 300 μl of sterile gel was added to the transwell insert and placed on a cell-seeded well and continued to culture at 37 °C with a 5 % CO_2_ incubator for three days. Biocompatibility was assessed every day for three days by using live/dead staining and cell proliferation assay.

To determine the viability of cells, live/dead staining would be performed upon indirectly culturing them with gels [33]. The transwell inserts were removed followed by withdrawing media and washed with 1X DPBS. Then, calcein AM (1 µM, #C3100MP, Invitrogen™) and ethidium homodimer (1 µM, #E1169, Invitrogen™) was added to the well and incubated at 37 °C with 5 % CO_2_ for 15 min. Then, fluorescently stained samples were washed with 1X DPBS and visualized under Keyence microscope (BZ-X series, Keyence microscope) under GFP channel for calcein AM (green color, viable cells) and ethidium homodimer at RFP channel (red color, dead cells). The proliferation of cells exposed to formulated gels was quantified using PrestoBlue™ cell viability reagent at different time points. In short, PrestoBlue™ (#A13262, Invitrogen™) working concentration solution (1:10 dilution with complete media) was prepared per the manufacturer’s instructions. Cell culture complete media was decanted from respective samples, followed by the addition of PrestoBlue™ solution and incubated for 3 h at 37 °C with 5 % CO_2_ incubator. After that, the solution was collected from each sample, and absorbance was measured at 570 nm and 600 nm with a Varioskan Lux plate reader (Thermo Fisher Scientific). With calculations from the absorbance data, it would be found that the cell proliferation rate is directly proportional to dye reduction percentage, which would be presented as a bar graph with standard deviation (triplicate samples were used for each time point) [34, 35].

To visualize the morphology of adhered cells following treatment with drug-loaded and non-loaded gels, actin filaments were stained using a fluorescent dye. After 3 days of culture, both HDFs and HaCaTs were fixed with 4% formaldehyde (#30450000, Bioworld) overnight at 4 °C, followed by washing with 1X DPBS. Cells were then permeabilized with 0.1% Triton X-100 for 10 minutes, washed three times with 1X DPBS, and blocked with 1% bovine serum albumin (#A3983-500G, Sigma-Aldrich) for 30 minutes. Alexa Fluor™ 594 phalloidin (#A12381, Fisher Scientific) was added to stain actin filaments, incubated at room temperature for 30 minutes, and then washed with 1X DPBS. Nuclei were stained in contrast for 5 minutes. Fluorescent samples were recorded via a Keyence BZ-X series microscope, capturing RFP (cytoskeleton, red) and DAPI (nuclei, blue) channels. Estimation of *in vitro* wound closure using scratch assay

*In vitro* wound closure experiments were performed using scratch assay on two different cell types [33]. Cells were then momentarily seeded on a 24-well plate at a seeding density of 5 × 10^4^ cells per well and cultured till they attained 70% confluency then using a sterile 10 µl tip, a direct line of scratch was made on attached cells to create breach between compact cell monolayer. Gel filled transwell inserts were placed in the well and supplemented with DMEM complete media to check the impact of degraded material/subproducts on cell survival. Bright-field images were captured using the Keyence microscope (BZ-X series, Keyence) at different time points such as 0, 24, and 48 h to compare the cell migration between sample groups. Collected images were quantified for the rate of cell migration and wound closure percentage using below mentioned formula with ImageJ software,

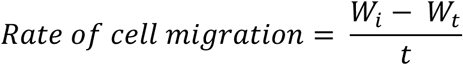

Where W_i_ is the initial wound width average in µm, W_t_ is wound width at different time points, and t is the time point of the assay in hours.

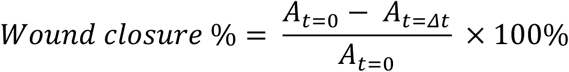

In this case, *A*_*t*=0_ refers to the initial wound area, and *A*_*t*=*Δt*_ is the wound area at an indicated time point after a time span of initial hours, both in µm^2^.

### *In vivo* mice burn wound model

#### Mice skin burn injury

Male C57BL/6J mice (10 weeks old, 23–28 g) were obtained from Jackson Laboratories (Bar Harbor, ME). All procedures were approved by the University of Arizona IACUC (Protocol #2023-1071). A severe third-degree burn wound was created following our established protocol [16]. In brief, the mice were anesthetized intraperitoneally (IP) using a cocktail of ketamine hydrochloride at a dose of 100 mg/kg and xylazine at 10 mg/kg, both sourced from Dechra Veterinary Products in Kansas, USA. A trimmer and depilatory cream (Nads) would be used to remove hair from the thoracolumbar dorsal region to create a burn injury. The target region was prepared for the burn injury using a 70% alcohol and a 5% povidone solution. A burn was created using a carbon rod with a surface area of 10 mm and weighing 65 g, with no external pressure. First, the rod was heated using an aluminum heating block at 95 °C and then applied on the skin for 4 s (**Fig. 4A**). After the creation of the burn wound, all animals received a buprenorphine (3.25 mg/kg, #NDC86084-100-30) as an analgesic to relieve pain. The mice in the burn wound experimental groups (n = 6 mice/ group) were chosen at random to test the control gel (gel without 4-AP) and 4-AP gel (5 µg of 4-AP in 500 mg gel) groups. The control and 4-AP gels (500 mg/mouse) were applied topically to the wound immediately after creating the burn and again on post-surgery dressing days 3, 6, 9, 12, 15, 18, and 21, every three days alternately. The wound area and test materials were covered with Tegaderm (#1610, 3M Health care). On day 21, the mice were euthanized, and their skin wound tissues were harvested and processed for experimental analyses.

#### Burn wound closure measurements

Post-burn wounds were captured using a digital camera on days 3, 6, 9, 12, 15, 18, and 21 (**Fig. 1A**). (**Fig. 1A**). Wound measurements were obtained through NIH ImageJ software (version 1.53e) from the National Institutes of Health (Bethesda, MD, USA), with a reference scale for accuracy. The % wound closure was calculated based on the measurements taken on day 0, using the formula below:

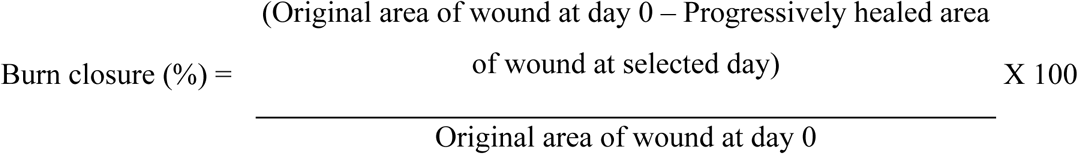

**Fig. 1.**
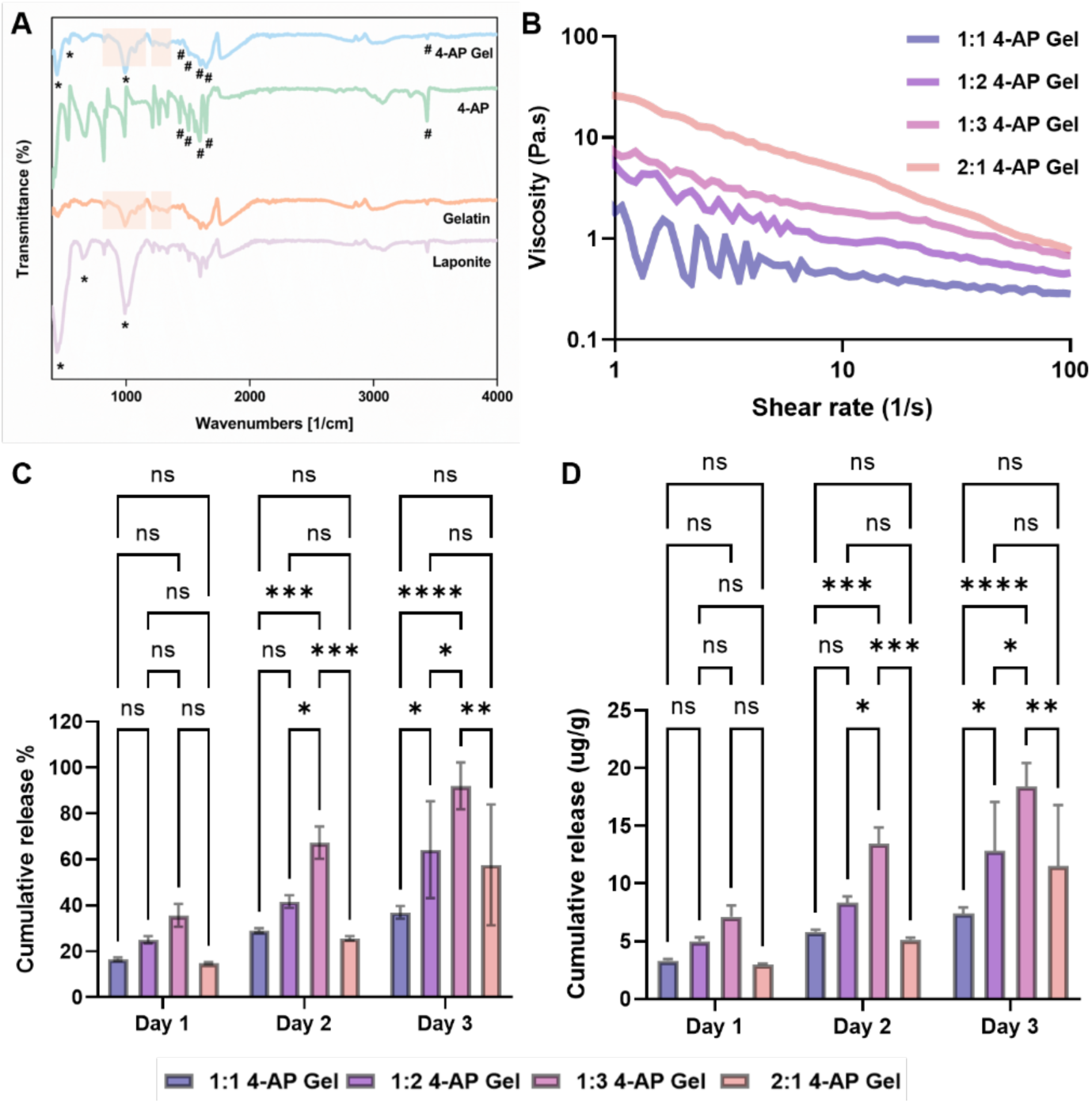
Formulation of 4-AP loaded laponite-gelatin gels and their characterizations. (**A**) Fourier transform infrared spectroscopy (FTIR) spectra, “*” denotes Laponite peaks while “#” for 4-AP spectra. (**B**) Viscosity profile measurements performed at constant room temperature. (**C**) Cumulative drug release percentage from different formulations. (**D**) The release of 4-AP drug (µg) per g of gel at different time points with four different gel compositions. * p<0.05, ** p<0.01, ***p<0.001, ****p<0.0001 and ns for non-significant where p>0.05.

### Histological staining

The analysis of skin tissue hematoxylin and eosin (H&E) staining followed our established protocol [16]. In brief, skin samples were collected from the area surrounding the target region using a 12 mm acu-punch (#NC9253254, Fisher Scientific). The tissues were evenly divided at the wound’s center for histological, gene, and protein expression analyses. The skin tissues were submerged in a 4% paraformaldehyde solution (#SC281692, ChemCruz) for 12 to 14 h and then washed three times with 70% alcohol before being embedded in paraffin. Each section was cut to a thickness of 5 μm from the tissue blocks (#HM315, GMI). The paraffin was removed while the tissue was gradually rehydrated using xylene and alcohol and then stained with an H&E kit (#ab245880, Sigma-Aldrich). First, sections would be stained using modified Mayer’s hematoxylin for 5 min, followed by a 15 s incubation with a bluing reagent. Then, they were stained with eosin (#71204, Thermo Scientific) for 20 s and dehydrated using 95% and 100% alcohol for 5 min each, twice. Next, the sections were fixed in xylene for 5 min, twice, and finally mounted with DPX (#06522, Sigma-Aldrich). The prepped slides were scanned at 80x magnification by use of a slide scanner (MoticEasyScan, SF, USA). The wound length, wound area, epithelial tongue length, and number of hair follicles were quantified using NIH ImageJ-1.53e software.

### Immunofluorescence (IF) staining and analysis

IF staining of skin tissues were performed following our previously established method (39366954). In brief, antigen unmasking was performed using a 10 mM sodium citrate buffer (pH 6.0) at 95 °C for 20 min. Next, the tissues were permeabilized, and nonspecific binding was blocked with 1% Triton X-100 and 5% goat serum. The following antibodies were applied by incubating the slides overnight at 4 °C: K14 (1:100, # NBP2-34270, Novus Biologicals), K15 (1:500, #833901, BioLegend), Vimentin (1:200, # 10366-1-AP, Thermo Fisher Scientific), α-SMA (1:500, #14-9760-82, Thermo Fisher Scientific), and TGF-β (1:500, # 3711s, Cell Signaling). Next, the slides were rinsed three times in PBS, followed by incubation with the appropriate secondary antibodies: Alexa Fluor 488 (1:1000, #A11008, Invitrogen), Alexa Fluor 594 (1:1000, #A11032, Invitrogen) and Alexa Fluor 647 (1:1000, #A21247, Invitrogen) for 1 h at room temperature. Anti-fade reagent with DAPI (# P36935, Thermo Fisher Scientific) would be utilized to stain the nuclei. Finally, the slides were examined and recorded with a fluorescence microscope (#DM6000, Leica, IL, USA), and the following images were analyzed and quantified using NIH ImageJ version 1.53e software.

### Collagen staining and examination

Collagen staining of skin tissue was carried out using Herovici’s staining protocol (#KTHER, StatLab). Slides were first deparaffinized, then immersed in Weigert’s hematoxylin for 5 minutes. Next, the slides were rinsed with water for 45 s and then stained with Herovici’s solution for 2 min. After a 2-min wash in 1% acetic acid, the tissues were dehydrated in 100% alcohol and cleared in xylene, with each step repeated three times for 1 min. Lastly, an organic mounting medium (#06522, Sigma-Aldrich) was utilized to mount the slides, which were then imaged at 80× magnification by a MoticEasyScan (SF, USA). Collagen types I and III expression was quantified using NIH ImageJ software (version 1.53e).

### Gene expression analysis

RNA was extracted from skin tissue (#74104, Qiagen) and reverse-transcribed into cDNA using 1000 ng of RNA (#4368814, Applied Biosystems). qRT-PCR was performed with gene-specific primers (Table S1) and qPCR Master Mix (#4367659, Applied Biosystems) on an Azure Cielo 6 Real-Time PCR System. Relative mRNA expression levels were normalized to GAPDH, and fold changes were calculated by comparing the 4-AP gel group to the control.

### Protein expression analysis

Protein extraction from skin wound tissues and subsequent Western blot analysis were conducted following our previously established protocols [16]. Briefly, proteins were extracted from skin samples using extraction reagent (#78510, Thermo Fisher Scientific). Tissue homogenization was performed at 4 °C using stainless steel beads (#SSB14B, Next Advance) with a Bullet Blender homogenizer (#BBX24, Next Advance), initially set at 5,000 rpm for 5 min, before being switched to 13,000 rpm for 20 min. Extracted proteins were then quantified with the BCA Protein Assay Kit (#23225, Thermo Fisher Scientific)For protein detection, 50 μg of total protein was separated using 4–12% SDS-PAGE (#4561044, GenScript). Following electrophoresis, proteins were transferred to a membrane (#L00686, GenScript) and blocked with 3% BSA in 1X TBST for 1 hour at 37°C. Next, the membranes were incubated with the primary antibodies listed below for 12 to 14 h, at 4°C: K14 (1:3000, #NBP2-34270, Novus Biologicals), K15 (1:1000, #833901, BioLegend), Vimentin (1:5000, #10366-1-AP, Thermo Fisher Scientific), α-SMA (1:500, #14-9760-82, Thermo Fisher Scientific), TGF-β (1:1000, #3711S, Cell Signaling), and β-Actin (1:5000, #A1978, Sigma). After three washes with TBST, membranes were incubated with either anti-rabbit HRP-conjugated (#7074, Cell Signaling) or anti-mouse HRP-conjugated (#7076, Cell Signaling). Signal detection was conducted using chemiluminescent substrate (#34579, Thermo Fisher Scientific), and bands were visualized with a G:BOX Chemi XRQ gel imaging system. Densitometric analysis was conducted using NIH ImageJ software (version 1.53e). Full, unedited Western blot images of all probed proteins are provided in the supplemental materials.

### Statistical analysis

All experiments were conducted using triplicate samples (n = 3), and data are presented as mean ± standard deviation (SD). Statistical analyses for in vitro studies were performed using GraphPad Prism 9. One-way ANOVA followed by Tukey’s post-hoc test was employed to assess differences among groups. Statistical significance was defined as follows: *p < 0.05, **p < 0.01, ***p < 0.001, ****p < 0.0001, and ‘ns’ for non-significant results (p > 0.05). *In vivo* studies comparing two groups (n ≥ 3) were analyzed using two-tailed, unpaired t-tests with GraphPad Prism 10.1.1. Results are presented as mean ± SEM, with significance levels indicated as follows: *p < 0.05, **p < 0.0021, ***p < 0.0002, and ****p < 0.0001.

## Results

### 4-AP loaded gel formulation and characterization

Injectable 4-AP drug loaded gel was formulated with varied combinations of laponite and gelatin. The functional groups of the samples were identified using FTIR spectroscopy. The major peaks corresponding to laponite, gelatin, and 4-AP were clearly observed in the drug-loaded laponite-gelatin sample, indicating the successful incorporation of the drug into the gel (**Fig. 1A**). The characteristic peaks of laponite—460 cm⁻¹ for Si–O–Mg, 652 cm⁻¹ for Mg–OH–Mg, and 1000 cm⁻¹ for Si–O bending vibrations—were present in both pristine laponite and the 4-AP-loaded gel [27]. Gelatin exhibited Amide II and III bands in the range of 670–1240 cm⁻¹ and 1400–1550 cm⁻¹ [29, 36]. The N–H asymmetric stretching vibration and in-plane deformation of 4-AP were observed around 3436 cm⁻¹ and 1648 cm⁻¹, respectively. The C=N stretching vibration of the pyridine ring appeared at 1602 cm⁻¹, while C=C stretching vibrations of the pyridine ring were identified near 1436 cm⁻¹ and 1507 cm⁻¹ [37]. Primarily, different combinations of laponite and gelatin gels (**Table 1**) were formulated to identify their gel stability characteristics. Formulations started with 1:1 ratio of laponite to gelatin with increasing up to 1:3 then inversed laponite to gelatin 2:1 to check how changing these combinations influence the gel characteristics and drug release profile. The injectable properties of the formulated gels were evaluated through viscosity profile measurements. The viscosity vs. shear rate graph (**Fig. 1B**) demonstrated shear-thinning behavior for all samples, where an initially higher viscosity indicates stability at rest, followed by a decrease in viscosity under higher shear rates. This suggests that, upon external load or pressure, the gels can flow. Moreover, amplitude sweep tests confirmed the viscoelastic behavior of the gels, where the storage modulus (G’) was higher than the loss modulus (G”) in the 1:3 and 2:1 sample, indicating the most stable gels in all four combinations as shown in **Fig. S1A and B**. The frequency sweep test corroborated the amplitude results, with higher gelatin and laponite ratios (such as 1:3 and 2:1) showing stable gels, where the storage modulus (G’) maintained a plateau over a wide frequency range (**Fig. S1C and D)**, indicating a strong elastic network capable of energy storage.

To further investigate the drug release properties, initial drug release experiments were conducted using four different compositions of the laponite-gelatin gel to understand the release kinetics in relation to varying ratios of laponite and gelatin. The cumulative drug release percentage graph (**Fig. 1C**) revealed that the 1:3 4-AP gel exhibited the most significant release of 4-AP compared to the other formulations, it was observed that increasing gelatin composition (1:2 and 1:3) in the gel enhances the drug release and higher concentration of laponite (2:1) slows down the release profile due to stiff gel network limiting slow molecule movement/migration in the gel system. Additionally, the cumulative drug release per day graph (**Fig. 1D**) depicted a gradual and sustained release of 4-AP from the 1:3 4-AP gel, with approximately 20 µg/g of the drug released over a period of 3 days. From **Fig. S2**, it was observed that at least 5 µg of 4-AP has been released from 1:3 4-AP Gel which is higher than other sample groups.

### 4-AP loaded gels foster cell growth and proliferation

To evaluate cell viability upon interaction with drug-loaded gels, live/dead staining was performed at different time points. **Fig. 2A and C** show the viability of HDF and HaCaT cells, with intense green fluorescence staining indicating high cell viability and negligible dead cells (red color) and continued to proliferate over time. Additionally, it was observed that the cells formed a complete cytoskeleton, with cell morphology assessed using actin staining, as shown in **Fig. S3A and B**. To further confirm cell proliferation, a PrestoBlue™ assay was performed. The percentage of dye reduction directly correlates with cell proliferation. Increased metabolic activity was observed over time in all sample groups, indicating that the 4-AP Gel did not compromise cell growth when compared with the control samples (**Fig. 2B and D**).

**Fig. 2.**
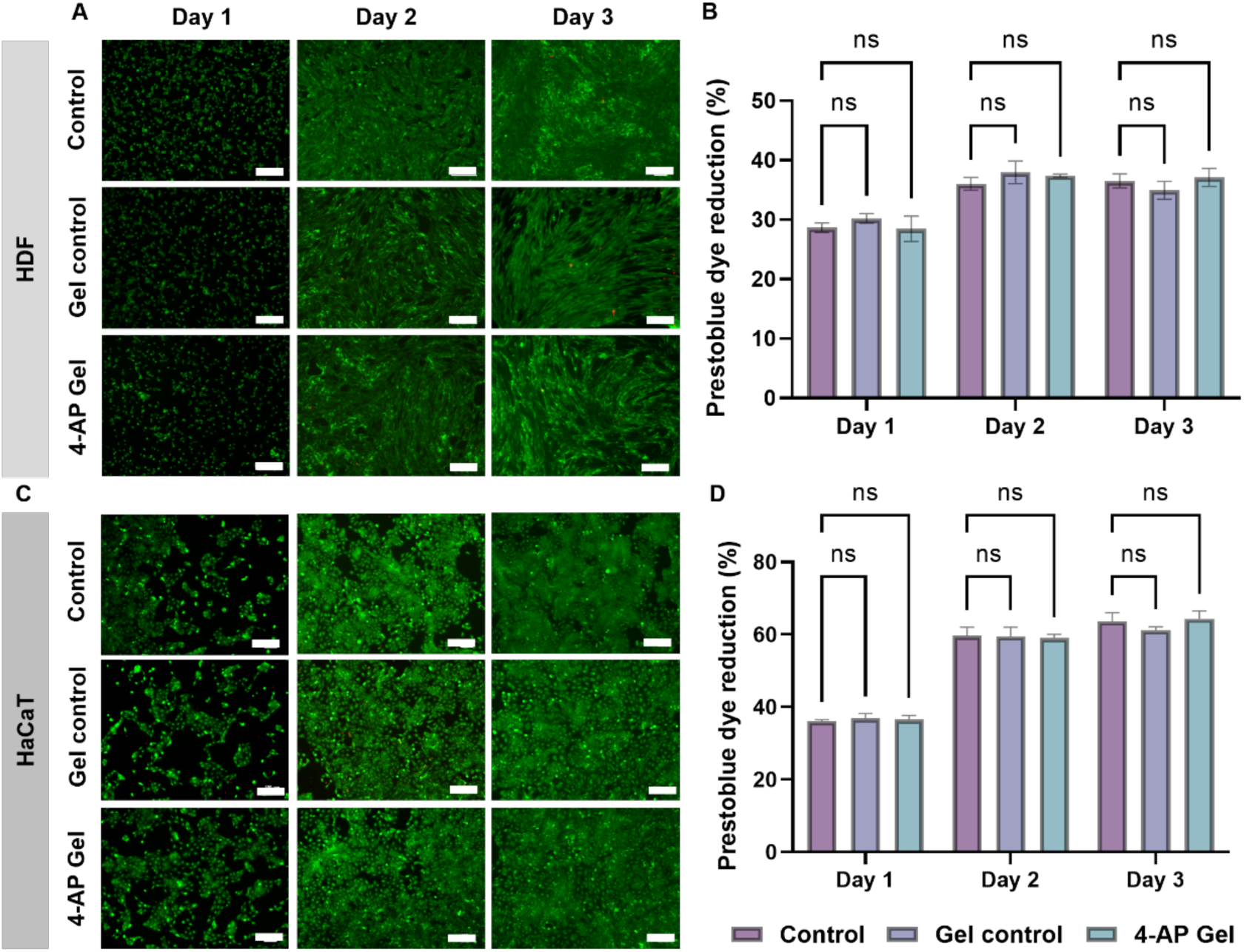
*In vitro* biocompatibility of human dermal fibroblasts (HDF) and human keratinocytes (HaCaT) upon interaction with 4-AP loaded gels. (**A and C**) Live/dead staining images after day 1, 2 and 3 days of culture with drug-loaded gels, green fluorescence denotes live cells while red represents dead cells (Scale bar 200 µm). (**B and D**) PrestoBlue™ assay showing reduction of dye over time, correlating cell proliferation. * p<0.05, ** p<0.01, ***p<0.001, ****p<0.0001 and ns for non-significant where p>0.05.

### 4-AP loaded gels supports cell migration in *in vitro* scratch assay

An *in vitro* scratch assay was conducted to evaluate the cell migration capacity of 4-AP gels and compare it with Gel control and the Control group using two different cell types. **Fig. 3A** shows that the migration of HDFs was unaffected by the presence of laponite-gelatin and 4-AP gels compared to the control group. Furthermore, the migration rate and wound closure percentage were calculated from the captured images, which indicated that the drug-loaded gel did not negatively affect migration (**Fig. 3B and C**).

**Fig. 3.**
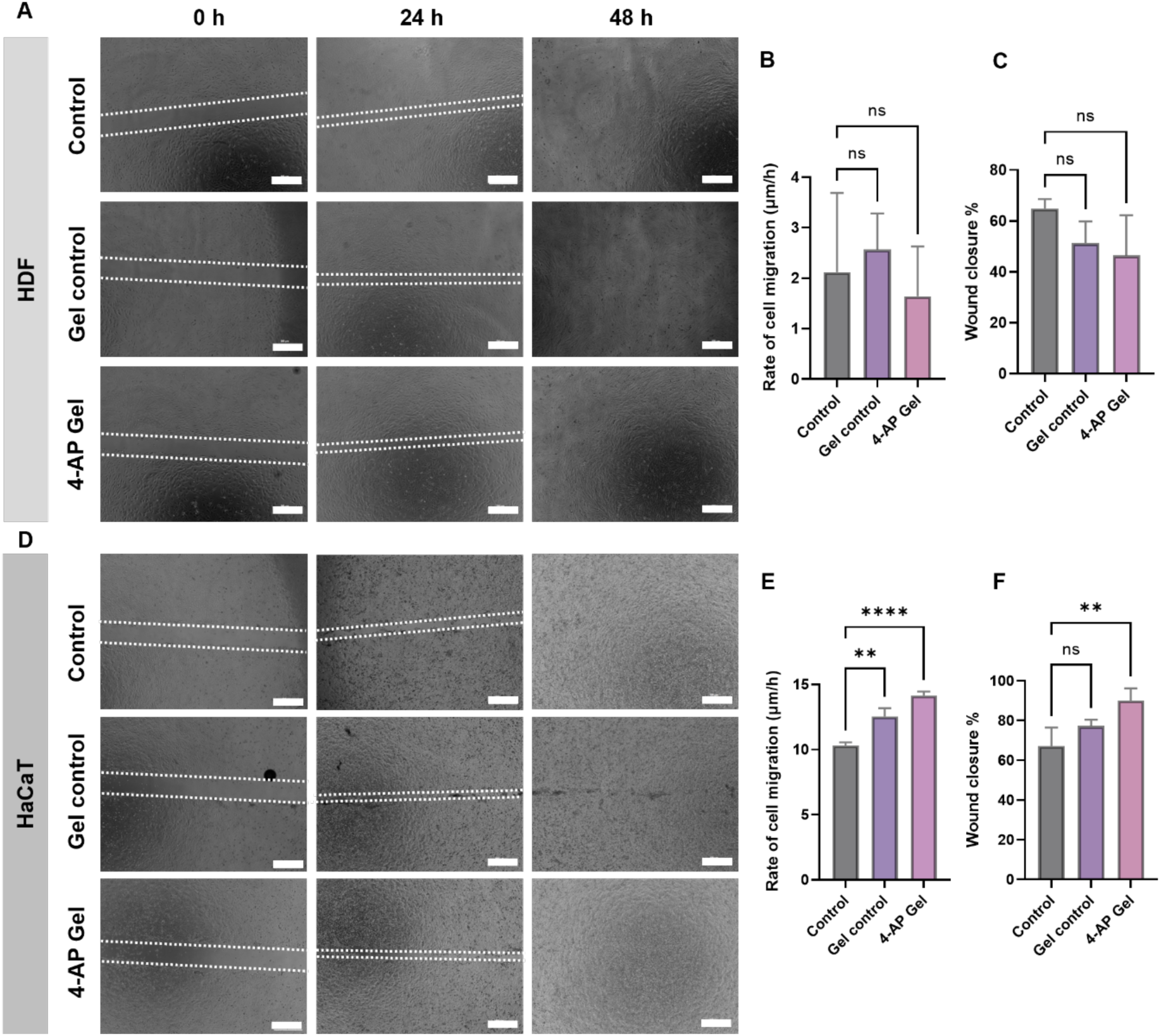
*In vitro* scratch assay performed on HDF and HaCaT cells. (**A and D**) Representative brightfield microscopy images of synthetic wounds created on monolayer cells, with cell migration observed over time to cover the scratch area (scale bar: 500 µm). (**B and C**) Rate of cell migration and wound closure percentage in HDFs, calculated from captured images using ImageJ. (**E and F**) Rate of cell migration and wound closure percentage in HaCaT cells, calculated from captured images using ImageJ. * p<0.05, ** p<0.01, ***p<0.001, ****p<0.0001 and ns for non-significant where p>0.05.

Additionally, wound healing studies were conducted on keratinocytes, a major cell type in the skin. **Fig. 3D** shows 100% wound closure in all groups within 48 hours which corroborates HDF scratch assay results (**Fig. 3A**). The rate of cell migration and wound closure was notably quicker in the 4-AP gel group compared to the controls, as demonstrated in **Fig. 3E and F**. To further investigate wound repair, *in vivo* experiments were conducted using a burn wound mouse model.

### 4-AP expedited wound closure and regeneration following skin burn

We hypothesize that the topical application of 4-AP gel accelerates burn wound closure and promotes tissue regeneration in severe third-degree burns, where we have previously observed the benefits of systemic 4-AP administration in a mouse model [16]. The details of our approved burn model are illustrated in **Fig. 4A**. We used macroscopic images of skin burns (**Fig. 4B**) to measure the percentage of wound closure over time (**Fig. 4C**). On day 3, mice treated with 4-AP gel demonstrated only a 5.50 % expansion in their original burn area, while those treated with the control gel showed an average expansion of 18.92 % (*P < 0.05). This significant impact can be endorsed to promote early protection against the common secondary expansion exhibited byof burn injuries. Burn closure in the 4-AP gel group also consistently outperformed versus control gel group on day 6 (30.24 % vs. 10.91 %), day 9 (50.32 % vs. 29.16 %), day 12 (70.51 % vs. 50.83 %), day 15 (79.34 % vs. 61.55 %), day 18 (89.16 % vs. 77.21 %), and day 21 (94.21 % vs. 81.87 %). Onday 21, mice tested with 4-AP showed nearly complete healing, while those given the control gel did not. Our results indicate that the application of 4-AP gel significantly accelerates burn wound closure versus the control gel (*P < 0.05, **P < 0.0021, and ***P < 0.0002).

**Fig. 4.**
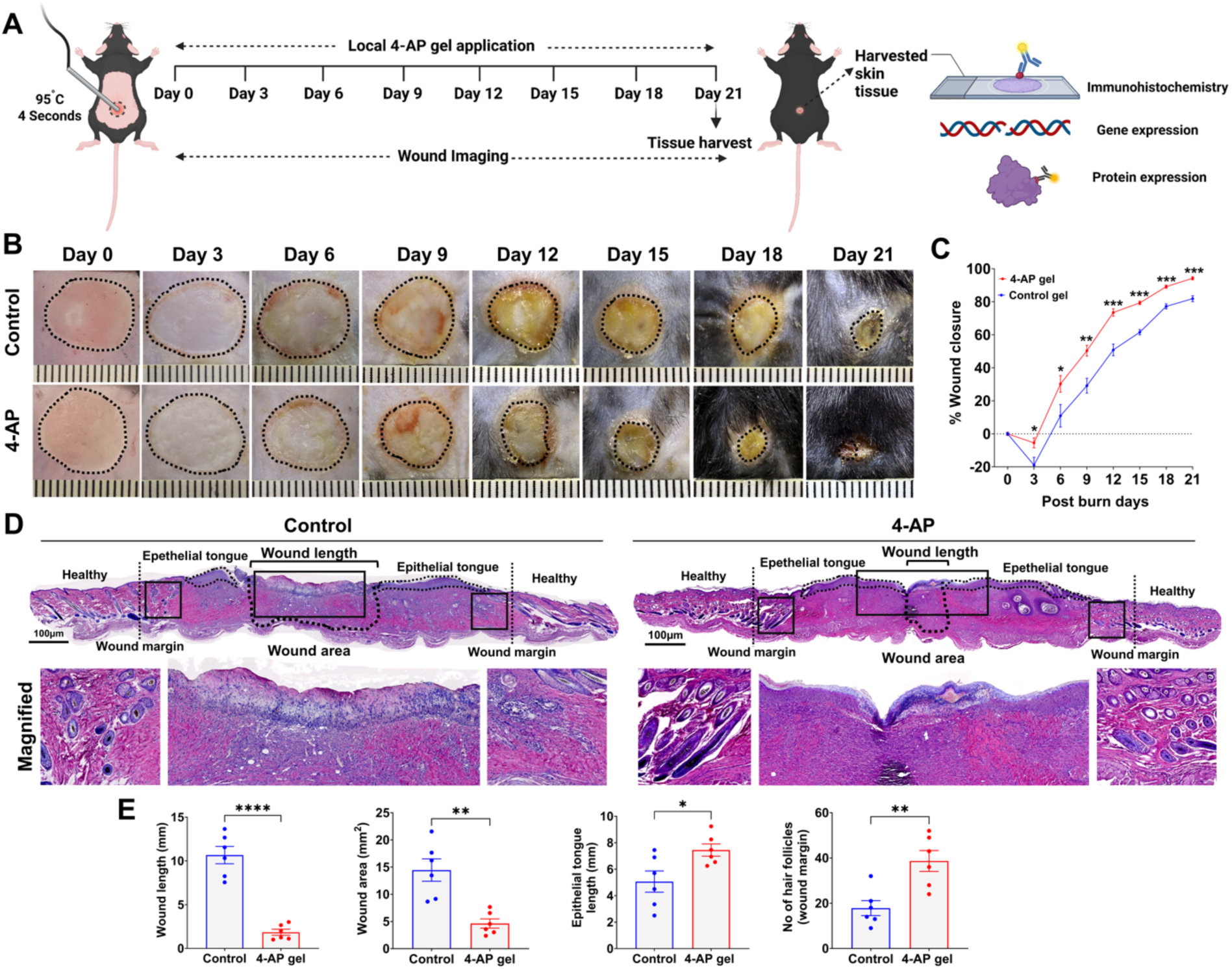
4-AP gel accelerated burn wound closure and improved skin regeneration. **(A)** Visual representation of study details to evaluate effects of the 4-AP gel in a C57BL/6 skin burn mouse subject. **(B)** Representative macroscopic records showing the healing of skin burn wounds treated with control and 4-AP gels (5 µg of 4-AP in 500 mg gel) treated mice at 3, 6, 9, 12, 15, 18, and 21 days, every three days alternatively. Scale bar = 15 mm. **(C)** The percentage of progressive burn wound closure per each day, compared between control and 4-AP gel-treated mice. n = 6 animals per group. **D)** Images characterizing full-thickness skin burns stained with H&E, comparing control and 4-AP gel treatments on day 21. Scale bar = 100 µm. n = 6 skin tissue samples per group. **(E)** Quantitative results of burn wound length, wound size, epithelial tongue length, and the number of regenerating hair follicles were analyzed using H&E-stained sections of skin burns with ImageJ software on day 21. n = 6 mouse skin tissue samples per group. Data are presented as mean ± SEM. Statistical significance was determined with *P < 0.05, **P < 0.0021, ***P < 0.0002, and ****P < 0.0001 using two-tailed, unpaired t-tests. to the control gel group). Comparisons were made using two-tailed, unpaired t-tests.

We also measured wound area, wound length (the distance between the healing epidermal tongues), epithelial tongue length, and hair follicle regeneration in the wounded area using H&E histology (**Fig. 4D**). Results of burn wound length between control and 4-AP gel treated mice (**Fig. 4E**; 1.84 mm vs. 10.66 mm; *P < 0.05), burn wound area (**Fig. 4E**; 4.62 mm^2^ vs. 14.42 mm^2^; *P < 0.05), epithelial tongue length (**Fig. 1E**; 7.44 mm vs. 5.06 mm; *P < 0.05), and hair follicle regeneration (**Fig. 4E**; 38 vs. 17; *P < 0.05) at day 21 post-burn injury confirmed the accelerated healing effect of 4-AP gel compared to the control gel. Granulation is a crucial component of the wound-healing process, and the application of 4-AP gel significantly enhanced the formation of granulation tissue (pink to red coloration), confirming its role in wound regeneration.

### 4-AP accelerated re-epithelization and hair follicle regeneration following skin burn

Studies have shown that K14 and K15 are important markers of basal keratinocytes and stem cells that play an important role in the regeneration of the epidermis and hair follicles [38–40]. Given our previous findings that systemic 4-AP enhanced the regeneration of epidermal keratinocytes and dermal hair follicle stem cells after excision [15] and burn [16] wounds, we aimed to demonstrate the impact of these findings using a topical 4-AP gel for treating skin burns.

IF staining analysis of keratin markers K14 and K15 showed that the application of the 4-AP gel significantly increased their expression in both epidermis and hair follicles on day 21 following the skin burn injury, compared to the control gel group (**Fig. 5A and B**; 5.9 x 10^7^ vs. 3.2 x 10^7^; 6.6 x 10^7^ vs. 3.7 x 10^7^; **P < 0.0021). Representative tile scanning IF images of K14 and K15 are presented in **Fig. S4**. We also assessed the expression of K14 and K15 in burned skin tissues on day 21 and found that applying 4-AP gel significantly increased both protein (**Fig. 5C and D**; **P < 0.0021) and gene expression levels (**Fig. 5E**; *P < 0.05 and **P < 0.0021). These findings strongly suggest that the 4-AP gel positively effects re-epithelialization and hair follicle regeneration, like the effects we previously observed with systemic 4-AP in standard excision and skin burn wound healing.

**Fig. 5.**
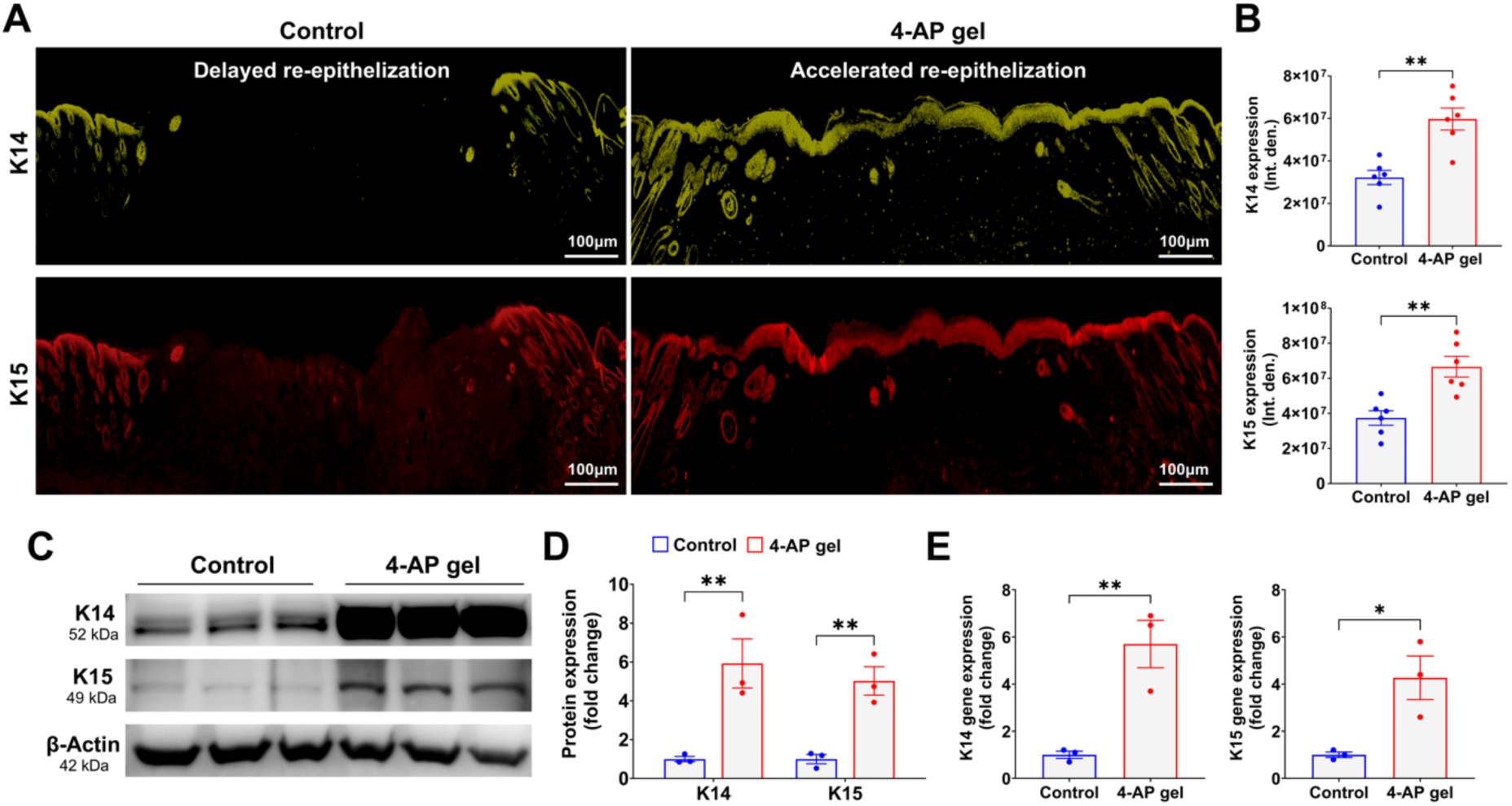
4-AP enhanced post-skin burn epithelialization and hair follicle regeneration. (A and. **B)** Representative IF images and quantitative results of keratinocyte and hair follicle regeneration markers (K14 and K15) in skin burn mice treated with control or 4-AP gels on day 21. Scale bar = 100 µm. n = 6 mouse skin tissue samples per group. Protein **(C and D)** and gene expression results **(E)** indicate that the 4-AP gel enhances the keratin markers K14 and K15 on day 21 post-skin burn injury, compared to the control group. Data from three skin tissue samples per group are shown as mean ± SEM, with significance marked (*P < 0.05 and **P < 0.0021). Comparisons were performed using two-tailed, unpaired t-tests.

### 4-AP increased fibroblast transformation after skin burn

Studies, including ours, have demonstrated that TGF-β is crucial for the transformation of fibroblasts to myofibroblasts, which is essential for ECM remodeling following skin burns [16, 41, 42]. The results from IF staining showed that 4-AP gel (vs. control gel) largely augmented the expression of vimentin and α-SMA in addition to TGF-β expression on day 21 post-skin burn (**Fig. 6A and B**; 6.3 x 10^7^ vs. 3.9 x 10^7^; 4.5 x 10^7^ vs. 2.8 x 10^7^; 4.8 x 10^7^ vs. 2.4 x 10^7^; *P < 0.05 and **P < 0.0021). Representative tile scanning IF images of vimentin, α-SMA, and TGF-β are presented in **Fig. S5**. We also investigated the effects of 4-AP gel on controlling gene and protein expression for these markers. Our findings revealed that 4-AP gel significantly elevated the levels of vimentin, α-SMA, and TGF-β (**Fig. 6C-E**; *P < 0.05 and **P < 0.0021) compared to the control gel treatment. These data strongly support the significance of 4-AP gel in controlling TGF-β downstream signaling during the remodeling of burn tissue.

**Fig. 6.**
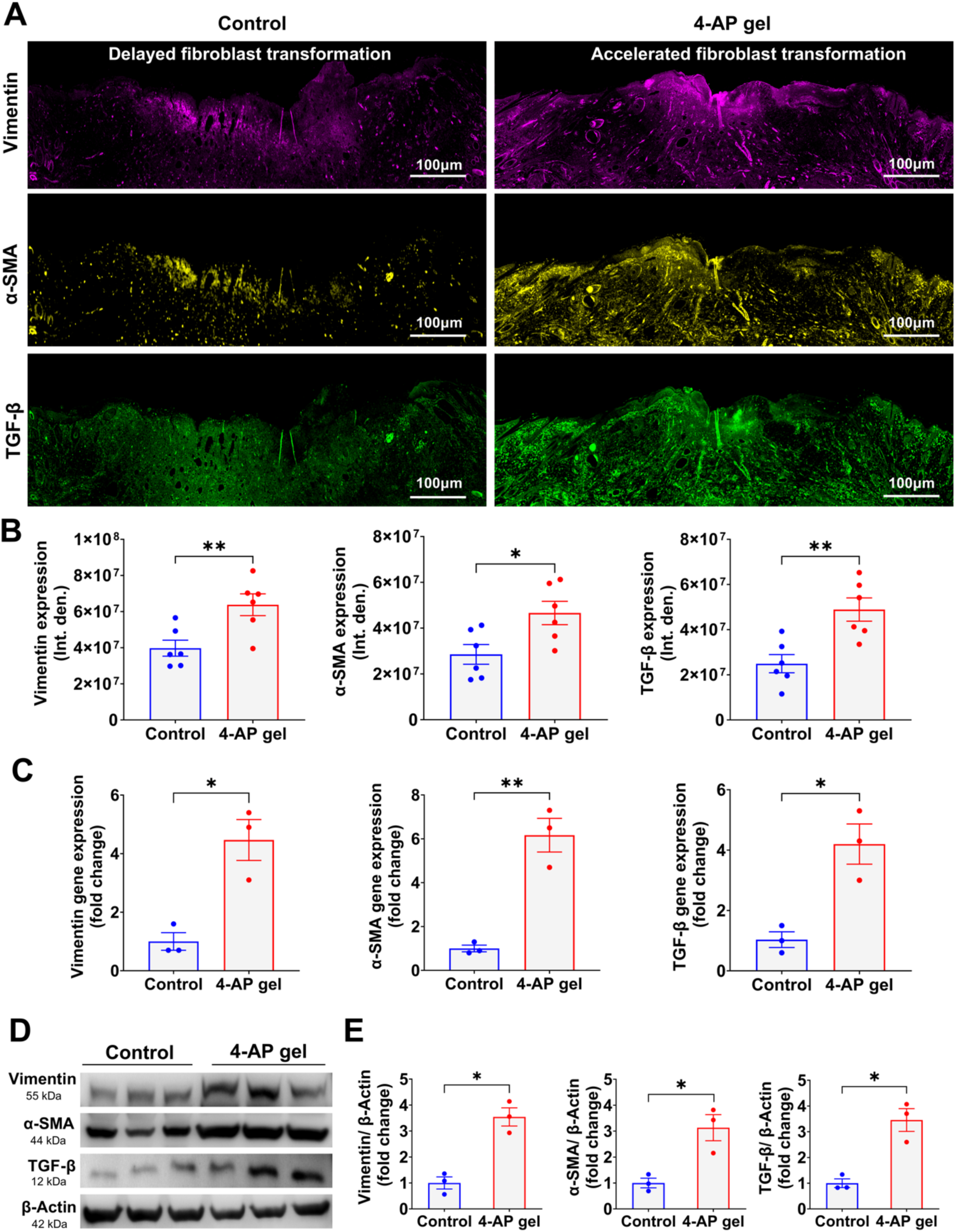
4-AP enhanced fibroblast transformation after skin burn. (A and. **B)** IF results of vimentin, α-SMA, and TGF-β in control mice vs. 4-AP gel after a skin burn injury on day 21 (scale bar = 100 µm, n = 6 skin tissue samples per group). **(C)** Gene expression results indicate that the 4-AP gel augmented vimentin, α-SMA, and TGF-β gene expression compared to the control gel group (n = 3 skin tissue samples per group). **(D and E)** 4-AP treatment considerably enhanced protein levels of these markers compared to the control gel group (n = 3 skin tissue samples per group). Data are presented as mean ± SEM, and comparisons were conducted using two-tailed, unpaired t-tests (*P < 0.05 and **P < 0.0021).

### 4-AP increased collagen matrix remodeling after skin burn

During the later stages of skin regeneration following a burn, fibroblasts and keratinocytes play critical roles in transformation of the ECM [43, 44]. Fibroblasts, particularly through their transition into myofibroblasts, produce ECM proteins such as collagen types I and III. These proteins, with their increased ratios, help improve the skin’s tensile strength and elasticity [45, 46]. On day 21 post-skin burn, we quantified the expression of collagen I and III using Herovici’s collagen staining technique. Our results demonstrated that the 4-AP gel significantly accelerated the percentages of collagen types I and III, and their ratios, compared to the control gel (**Fig. 7A and B**; 438 % vs. 267 %; 288 % vs. 215 %; 1.7 vs. 1.2; *P < 0.05, **P < 0.0021, and ***P < 0.0002). 4-AP gel application also significantly increased the gene expression of these markers compared to the control gel (**Fig. 7C**; **P < 0.0021 and ***P < 0.0002). These findings suggest that the 4-AP gel promotes remodeling of ECM during the healing process of burns. Schematic illustrations of skin burn healing and remodeling are shown in **Fig. 7D**.

**Fig. 7.**
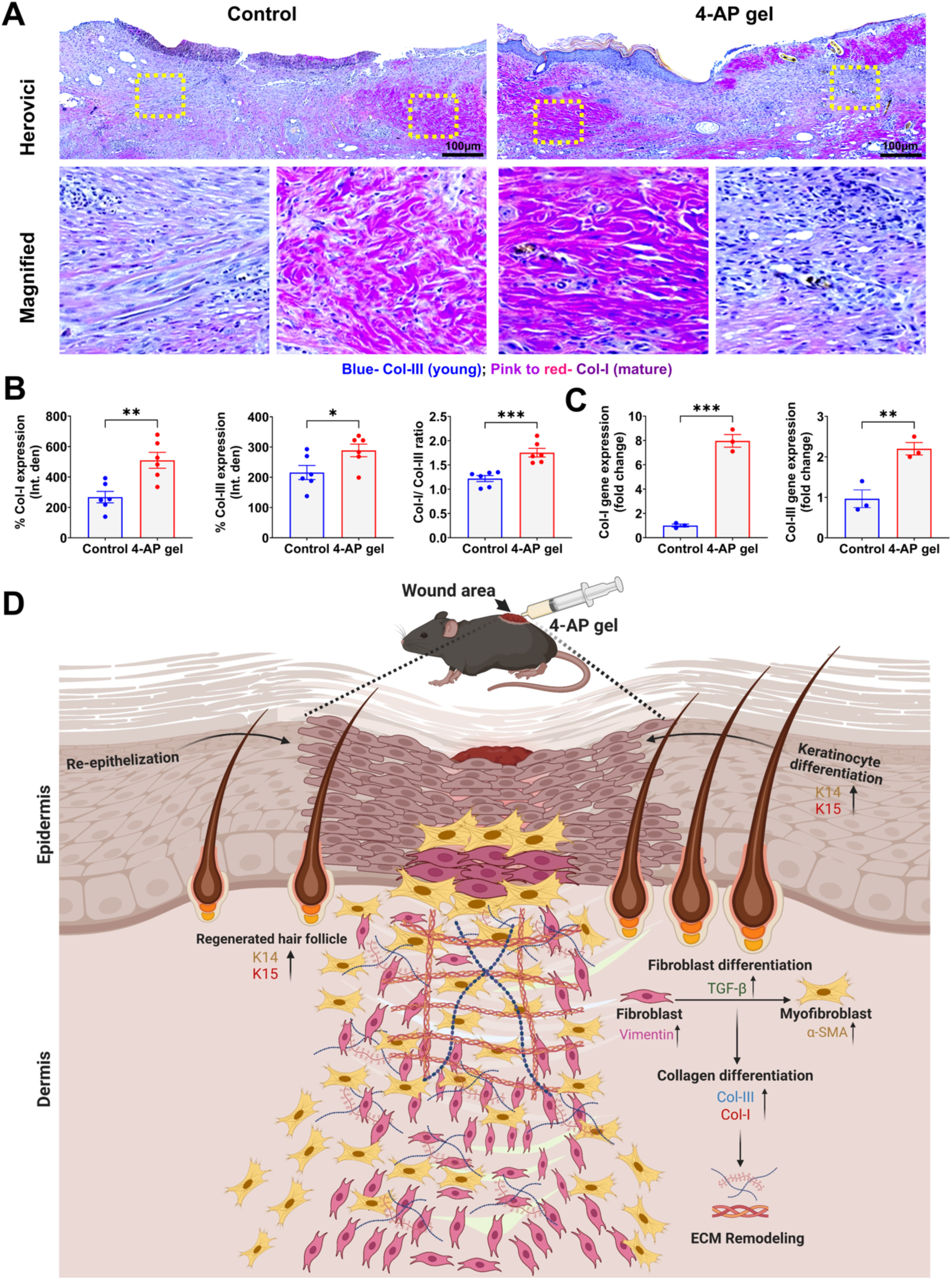
4-AP gel advanced skin burns extracellular matrix remodeling. (A and. **B)** Representative Herovici’s staining images and quantitative results of collagen-III (blue; younger collagen) and collagen-I (pink to red; mature collagen), as well as their ratios in the skin tissue samples from control and 4-AP gel-treated mice 21 days post-burn. n = 6 skin tissue samples per group. **(C)** 4-AP gel notably augmented collagen-III and collagen-I gene expressions versus the control gel group on day 21 post-burn skin injury. n = 3 skin tissue samples per group. Data are presented as the mean ± SEM, and comparisons were made using two-tailed unpaired t-tests (*P < 0.05, **P < 0.0021, and ***P < 0.0002). **(D)** The schematic illustration depicts the effects of 4-AP gel treatment on the healing process of skin burns. The 4-AP gel enhances the re-epithelialization and differentiation of keratinocytes and fibroblasts. This activity supports the remodeling of the ECM through collagen differentiation and TGF-β signaling.

## Discussion

Our previous studies demonstrated that the systemic administration of 4-aminopyridine (4-AP) accelerated the healing of both full-thickness excisional wounds and burn injuries by promoting re-epithelialization, dermal regeneration, and remodeling [15, 16]. The effects we observed involved multiple cell types, which led to obtaining exemptions from the US FDA to conduct clinical trials on excisional and burn wounds (ClinicalTrials.gov ID NCT06333171; NCT06596434; please cite these references as mentioned in the introduction). In the current study, we decided to investigate the unexplored subject of whether the topical application of a 4-AP gel, administered at lower doses every three days, could also accelerate the closure and regeneration of burn wounds, as opposed to using systemic daily high doses. Our results indicated that the 4-AP gel significantly enhanced both epidermal and dermal regeneration, thereby speeding up the closure of burn wounds.

Initially, different combinations of formulated gels were tested to study their characteristics. The results of the viscosity profile measurements highlight the shear-thinning behavior of the gels, which is a critical property for injectable formulations [27]. Notably, it was observed that increasing the concentrations of gelatin and laponite in the formulations led to an increase in the viscosity range, particularly in the 1:3 and 2:1 4-AP gels, which exhibited a stable shear-thinning profile (**Fig. 1b**). The initial high viscosity ensures the gel remains stable at rest, while the decrease in viscosity under shear suggests that the gels will flow when subjected to external pressure, facilitating their injection. This suggests that the gels could maintain their structural integrity during extrusion and exhibit ease of flow during application [47]. The amplitude sweep test demonstrated that the gels with higher gelatin and laponite ratios (1:3 and 2:1) exhibit superior stability, as evidenced by the storage modulus (G’) being consistently higher than the loss modulus (G”) in these formulations (**Fig. S1A and B**). This indicates that the gels possess a more robust and elastic network structure, making them more stable under mechanical stress. Additionally, the frequency sweep test (**Fig. S2A and B**) reinforced these findings, showing that these stable gels with higher gelatin and laponite ratios maintain a plateau in the storage modulus over a wide range of frequencies. This suggests that these gels can store energy, making them highly effective for their intended application in drug delivery systems. However, the gels also display the necessary flow properties upon deformation, ensuring they meet the functional requirements for injectable applications [48].

Drug release profile suggests that this particular composition such as 1:3 may offer the most effective drug release profile (**Fig. 1C and D**). This release rate is ideal for local drug delivery, as it ensures that the drug is released at a sufficiently fast pace to achieve therapeutic effects while also maintaining a controlled release over a few days. This controlled release profile is crucial for applications requiring steady drug delivery in a localized area, thus minimizing the need for frequent reapplication [49]. Biocompatibility results demonstrated that these gels did not affect cell growth and proliferation (**Fig. 2**). Moreover, synthetic *in vitro* wound model study demonstrated the 4-AP Gel enhances cell migration and wound closure in 48 h compared to Control group (**Fig. 3D-F**). These results suggest that the loaded drug enhances cell migration, potentially aiding the wound repair process.

After a burn injury, the skin’s response to heat is crucial for minimizing burn wound expansion and initiating tissue regeneration [16]. Burn wound healing is a multi-phased process that begins with hemostasis and inflammation, followed by proliferation and remodeling, leading to the terminal differentiation of cells [50, 51]. Keratinocytes, the primary cells in the epidermis, depend on fibroblasts for effective wound healing [52]. Fibroblasts, which show high vimentin expression, can also transform into myofibroblasts, characterized by high α-SMA expression, in response to keratinocyte signaling [53, 54]. This transformation depends on a carefully balanced early pro-inflammatory environment, followed by TGF-β signaling [52, 55]. Unfortunately, only a limited number of studies have investigated the complexities of the burn healing process with a local gel application, particularly focusing on the regeneration of both the epidermis and dermis to achieve effective wound closure [56, 57].

Our current study demonstrated that the local application of a 4-AP gel significantly accelerated wound closure from day 3 to day 21. Our H&E staining results supported these findings, showing tissue regeneration characterized by reduced wound length and area with regenerated hair follicles. We also conducted IF staining of K14 and K15 markers, where we noticed higher expression levels in the epidermis and dermal hair follicles of the 4-AP gel group compared to the control gel group. Studies have shown that K14 and K15 markers are highly expressed in proliferative and migratory keratinocytes, along with new hair follicle stem cells [58, 59]. In our previous study with full-thickness skin excisions, we noticed that systemic 4-AP significantly accelerated the expression of K14, K15, and K17 markers in in epidermal and dermal hair follicle regeneration [15]. This trend was also observed in severe third-degree skin burns, with increased expression of K10 and K15 [16]. Correspondingly, our current 4-AP gel findings support the hypothesis that it enhances the regeneration of both the epidermis and hair follicles. We believe these benefits may result from either the direct effects of 4-AP on keratinocytes or their neurogenic effects, which warrant further investigation of the subject in future studies.

After a burn, active immune cells recruit fibroblasts and keratinocytes to repair ECM proteins, including collagen types I and III, elastin, fibronectin, proteoglycans, and other related proteins [60, 61]. Notably, collagen I (70%) and collagen III (15%) form significant components of ECM proteins [62]. TGF-β is crucial for the proliferation and differentiation of fibroblasts, as indicated by high vimentin expression, into myofibroblasts characterized by elevated α-SMA expression [63]. We observed these effects in our prior publications with systemic 4-AP involving both excision and skin burn injuries [15, 16]. Our current data further corroborate these findings, demonstrating that the 4-AP gel enhances the expression of collagen I and III, along with their ratios at both gene and protein levels. This result strongly supports the functional role of 4-AP gel in ECM remodeling.

## Conclusion

Our study supports the use of 4-AP topical gel for treating severe burns and promoting skin regeneration versus systemic administration. This minimally invasive 4-AP gel enhances burn wound healing by promoting skin regrowth and remodeling underlying tissue through improved cell activation and differentiation. Our research encourages further investigation into how 4-AP interacts with immune cells, such as macrophages, and key cell types like keratinocytes and fibroblasts, which are essential for tissue regeneration. Furthermore, this local application could also be beneficial for treating atypical burn injuries like frostbites, radiation, chemical exposure, and electrical injuries, *etc*.

## Acknowledgement

The authors acknowledge The University of Arizona College of Medicine, Tucson, AZ, USA, for animal study experiments and Terasaki Institute for Biomedical Innovation, Los Angeles, CA, USA for supporting *in vitro* studies.

## CRediT authorship contribution statement

**Govindaraj Ellur:** Investigation, Data curation, Formal analysis, Methodology. **Meenakshi Kamaraj:** Investigation, Data curation, Formal analysis, Methodology, Writing – original draft, Writing – review and editing. **John C. Elfar**: Conceptualization, Investigation, Supervision, Funding acquisition, Writing – review and editing. **Johnson V. John:** Conceptualization, Investigation, Methodology, Supervision, Funding acquisition, Writing – review and editing. **Prem Kumar Govindappa:** Conceptualization, Investigation, Methodology, Supervision, Writing – original draft, Writing – review and editing. All authors read and approved the final manuscript.

## Data availability

The data presented in this study are available on request from the corresponding author.

## Ethical approval

Our manuscript does not contain any human data. Experimental design and animal protocols were approved by the Institutional Animal Care and Use Committee (IACUC) at The University of Arizona College of Medicine, Tucson, AZ. All experiments were conducted in accordance with the approved guidelines and regulations.

## Funding resources

This work was supported by grants from the National Institutes of Health (NIH; K08 AR060164-01A) and U.S. Department of Defense (DOD; W81XWH-16-1-0725) to JCE., NIH (R01DK134903) to JVJ, in addition to institutional support from The University of Arizona College of Medicine, Tucson, AZ, USA and Terasaki Institute for Biomedical Innovation, Los Angeles, CA, USA. The funding bodies played no role in the design of the study and collection, analysis, interpretation of data, and in writing the manuscript.

## References

[1] R.G. Frykberg, J. Banks, Challenges in the treatment of chronic wounds, Advances in wound care 4(9) (2015) 560–582.

[2] M.P. Rowan, L.C. Cancio, E.A. Elster, D.M. Burmeister, L.F. Rose, S. Natesan, R.K. Chan, R.J. Christy, K.K. Chung, Burn wound healing and treatment: review and advancements, Critical care 19 (2015) 1–12.

[3] N. Dasari, A. Jiang, A. Skochdopole, J. Chung, E.M. Reece, J. Vorstenbosch, S. Winocour, Updates in diabetic wound healing, inflammation, and scarring, Seminars in plastic surgery, Thieme Medical Publishers, Inc., 2021, pp. 153–158.

[4] P. Bowler, B. Duerden, D.G. Armstrong, Wound microbiology and associated approaches to wound management, Clinical microbiology reviews 14(2) (2001) 244–269.

[5] S. Nour, R. Imani, G.R. Chaudhry, A.M. Sharifi, Skin wound healing assisted by angiogenic targeted tissue engineering: A comprehensive review of bioengineered approaches, Journal of Biomedical Materials Research Part A 109(4) (2021) 453–478.

[6] S. Sarabahi, Recent advances in topical wound care, Indian journal of plastic surgery 45(02) (2012) 379–387.

[7] D. Snyder, N. Sullivan, D. Margolis, K. Schoelles, Skin substitutes for treating chronic wounds, (2020).

[8] A. Markiewicz-Gospodarek, M. Kozioł, M. Tobiasz, J. Baj, E. Radzikowska-Büchner, A. Przekora, Burn wound healing: clinical complications, medical care, treatment, and dressing types: the current state of knowledge for clinical practice, International journal of environmental research and public health 19(3) (2022) 1338.

[9] Z. Hussain, H.E. Thu, M. Rawas-Qalaji, M. Naseem, S. Khan, M. Sohail, Recent developments and advanced strategies for promoting burn wound healing, Journal of Drug Delivery Science and Technology 68 (2022) 103092.

[10] Y. Yao, A. Zhang, C. Yuan, X. Chen, Y. Liu, Recent trends on burn wound care: Hydrogel dressings and scaffolds, Biomaterials science 9(13) (2021) 4523–4540.

[11] A. Noor, A. Afzal, R. Masood, Z. Khaliq, S. Ahmad, F. Ahmad, M.-B. Qadir, M. Irfan, Dressings for burn wound: a review, Journal of Materials Science 57(12) (2022) 6536–6572.

[12] M.P. Sikka, J.A. Bargir, S. Garg, Modern developments in burn wound dressing, Research Journal of Textile and Apparel (2024).

[13] H.B. Jensen, M. Ravnborg, U. Dalgas, E. Stenager, 4-Aminopyridine for symptomatic treatment of multiple sclerosis: a systematic review, Therapeutic advances in neurological disorders 7(2) (2014) 97–113.

[14] F.A. Davis, D. Stefoski, J. Rush, Orally administered 4-aminopyridine improves clinical signs in multiple sclerosis, Annals of Neurology: Official Journal of the American Neurological Association and the Child Neurology Society 27(2) (1990) 186–192.

[15] M.G. Jagadeeshaprasad, P.K. Govindappa, A.M. Nelson, M.D. Noble, J.C. Elfar, 4-Aminopyridine induces nerve growth factor to improve skin wound healing and tissue regeneration, Biomedicines 10(7) (2022) 1649.

[16] R. VG, G. Ellur, A. A. Gaber, P.K. Govindappa, J.C. Elfar, 4-aminopyridine attenuates inflammation and apoptosis and increases angiogenesis to promote skin regeneration following a burn injury in mice, Cell Death Discovery 10(1) (2024) 428.

[17] T.C. Ezike, U.S. Okpala, U.L. Onoja, C.P. Nwike, E.C. Ezeako, O.J. Okpara, C.C. Okoroafor, S.C. Eze, O.L. Kalu, E.C. Odoh, Advances in drug delivery systems, challenges and future directions, Heliyon 9(6) (2023).

[18] M. Schäfer-Korting, W. Mehnert, H.-C. Korting, Lipid nanoparticles for improved topical application of drugs for skin diseases, Advanced drug delivery reviews 59(6) (2007) 427–443.

[19] S. Mishra, B. Reshma G, S. Pal, S. Bano, A. Gupta, A. Kumari, M. Ganguli, Topical Application of Peptide–Chondroitin Sulfate Nanoparticles Allows Efficient Photoprotection in Skin, ACS Applied Materials & Interfaces 13(2) (2021) 2382–2398.

[20] L. Zhao, J. Chen, B. Bai, G. Song, J. Zhang, H. Yu, S. Huang, Z. Wang, G. Lu, Topical drug delivery strategies for enhancing drug effectiveness by skin barriers, drug delivery systems and individualized dosing, Frontiers in Pharmacology 14 (2024) 1333986.

[21] D. Hawthorne, A. Pannala, S. Sandeman, A. Lloyd, Sustained and targeted delivery of hydrophilic drug compounds: A review of existing and novel technologies from bench to bedside, Journal of Drug Delivery Science and Technology 78 (2022) 103936.

[22] Y. Zhou, X.-L. Zhang, S.-T. Lu, N.-Y. Zhang, H.-J. Zhang, J. Zhang, J. Zhang, Human adipose-derived mesenchymal stem cells-derived exosomes encapsulated in pluronic F127 hydrogel promote wound healing and regeneration, Stem cell research & therapy 13(1) (2022) 407.

[23] F.S. Palumbo, M. Calligaris, L. Calzà, C. Fiorica, V.A. Baldassarro, A.P. Carreca, L. Lorenzini, A. Giuliani, C. Carcione, N. Cuscino, Topical application of a hyaluronic acid-based hydrogel integrated with secretome of human mesenchymal stromal cells for diabetic ulcer repair, Regenerative Therapy 26 (2024) 520–532.

[24] S. Uman, A. Dhand, J.A. Burdick, Recent advances in shear-thinning and self-healing hydrogels for biomedical applications, Journal of Applied Polymer Science 137(25) (2020) 48668.

[25] A. Sheikhi, S. Afewerki, R. Oklu, A.K. Gaharwar, A. Khademhosseini, Effect of ionic strength on shear-thinning nanoclay–polymer composite hydrogels, Biomaterials science 6(8) (2018) 2073–2083.

[26] N. Falcone, M. Ermis, A. Gangrade, A. Choroomi, P. Young, T.G. Mathes, M. Monirizad, F. Zehtabi, M. Mecwan, M. Rodriguez, Drug-Eluting Shear-Thinning Hydrogel for the Delivery of Chemo-and Immunotherapeutic Agents for the Treatment of Hepatocellular Carcinoma, Advanced Functional Materials 34(8) (2024) 2309069.

[27] N.R. de Barros, A. Gangrade, A. Rashad, R. Chen, F. Zehtabi, M. Ermis, N. Falcone, R. Haghniaz, S. Khosravi, A. Gomez, Injectable nanoengineered adhesive hydrogel for treating enterocutaneous fistulas, Acta biomaterialia 173 (2024) 231–246.

[28] F. Zehtabi, A. Gangrade, K. Tseng, R. Haghniaz, R. Abbasgholizadeh, H. Montazerian, D. Khorsandi, J. Bahari, A. Ahari, N. Mohaghegh, Injectable Shear-Thinning Hydrogels with Sclerosing and Matrix Metalloproteinase Modulatory Properties for the Treatment of Vascular Malformations, Advanced Functional Materials 33(51) (2023) 2305880.

[29] C. Li, C. Mu, W. Lin, T. Ngai, Gelatin effects on the physicochemical and hemocompatible properties of gelatin/PAAm/laponite nanocomposite hydrogels, ACS applied materials & interfaces 7(33) (2015) 18732–18741.

[30] J.W. Davern, L. Hipwood, L.J. Bray, C. Meinert, T.J. Klein, Addition of Laponite to gelatin methacryloyl bioinks improves the rheological properties and printability to create mechanically tailorable cell culture matrices, APL bioengineering 8(1) (2024).

[31] H. Takahashi, S. Ohnishi, Y. Yamamoto, T. Hayashi, N. Murao, M. Osawa, T. Maeda, K. Ishikawa, N. Sakamoto, E. Funayama, Topical application of conditioned medium from hypoxically cultured amnion-derived mesenchymal stem cells promotes wound healing in diabetic mice, Plastic and Reconstructive Surgery 147(6) (2021) 1342–1352.

[32] M. Kamaraj, A. Datla, S.E. Moulton, S.N. Rath, Biomimetic Mineralization of Mn-Doped Biphasic Calcium Phosphate in the GelMa Hydrogel Acting as a Functional 3D Bioscaffold for Osteo Defect Repair, ACS Applied Polymer Materials 6(1) (2023) 943–955.

[33] M. Kamaraj, N. Moghimi, A. McCarthy, J. Chen, S. Cao, A.R. Chethikkattuveli Salih, A. Joshi, V. Jucaud, A. Panayi, S.R. Shin, Granular Porous Nanofibrous Microspheres Enhance Cellular Infiltration for Diabetic Wound Healing, ACS nano (2024).

[34] M. Kamaraj, O. Rezayof, A. Barer, H. Kim, N. Moghimi, A. Joshi, M.R. Dokmeci, A. Khademhosseini, F. Alambeigi, J.V. John, Development of silk microfiber-reinforced bioink for muscle tissue engineering and in situ printing by a handheld 3D printer, Biomaterials Advances (2024) 214057.

[35] M. Kamaraj, U.K. Roopavath, P.S. Giri, N.K. Ponnusamy, S.N. Rath, Modulation of 3D printed calcium-deficient apatite constructs with varying Mn concentrations for osteochondral regeneration via endochondral differentiation, ACS Applied Materials & Interfaces 14(20) (2022) 23245–23259.

[36] S. Mohammadzadehmoghadam, Y. Dong, Fabrication and characterization of electrospun silk fibroin/gelatin scaffolds crosslinked with glutaraldehyde vapor, Frontiers in Materials 6 (2019) 91.

[37] Y. Chen, Y. Ma, Q. He, Q. Han, Q. Zhang, Q. Chen, Construction of pyridinium/N-chloramine polysiloxane on cellulose for synergistic biocidal application, Cellulose 26 (2019) 5033–5049.

[38] R. Yang, F. Liu, J. Wang, X. Chen, J. Xie, K. Xiong, Epidermal stem cells in wound healing and their clinical applications, Stem cell research & therapy 10 (2019) 1–14.

[39] Y. Guo, C.J. Redmond, K.A. Leacock, M.V. Brovkina, S. Ji, V. Jaskula-Ranga, P.A. Coulombe, Keratin 14-dependent disulfides regulate epidermal homeostasis and barrier function via 14-3-3σ and YAP1, Elife 9 (2020) e53165.

[40] K. Kretzschmar, F.M. Watt, Markers of epidermal stem cell subpopulations in adult mammalian skin, Cold Spring Harbor perspectives in medicine 4(10) (2014) a013631.

[41] M. Pakyari, A. Farrokhi, M.K. Maharlooei, A. Ghahary, Critical role of transforming growth factor beta in different phases of wound healing, Advances in wound care 2(5) (2013) 215–224.

[42] J.W. Penn, A.O. Grobbelaar, K.J. Rolfe, The role of the TGF-β family in wound healing, burns and scarring: a review, International journal of burns and trauma 2(1) (2012) 18.

[43] F. Cialdai, C. Risaliti, M. Monici, Role of fibroblasts in wound healing and tissue remodeling on Earth and in space, Frontiers in bioengineering and biotechnology 10 (2022) 958381.

[44] X. Dong, H. Xiang, J. Li, A. Hao, H. Wang, Y. Gou, A. Li, S. Rahaman, Y. Qiu, J. Li, Dermal fibroblast-derived extracellular matrix (ECM) synergizes with keratinocytes in promoting re-epithelization and scarless healing of skin wounds: Towards optimized skin tissue engineering, Bioactive Materials 47 (2025) 1–17.

[45] D. Singh, V. Rai, D.K. Agrawal, Regulation of collagen I and collagen III in tissue injury and regeneration, Cardiology and cardiovascular medicine 7(1) (2023) 5.

[46] L. Li, Y. Ma, G. He, S. Ma, Y. Wang, Y. Sun, Pilose antler extract restores type I and III collagen to accelerate wound healing, Biomedicine & Pharmacotherapy 161 (2023) 114510.

[47] H. Herrada-Manchón, M.A. Fernández, E. Aguilar, Essential guide to hydrogel rheology in extrusion 3D printing: how to measure it and why it matters?, Gels 9(7) (2023) 517.

[48] S. Jacob, A.B. Nair, J. Shah, N. Sreeharsha, S. Gupta, P. Shinu, Emerging role of hydrogels in drug delivery systems, tissue engineering and wound management, Pharmaceutics 13(3) (2021) 357.

[49] S. Adepu, S. Ramakrishna, Controlled drug delivery systems: current status and future directions, Molecules 26(19) (2021) 5905.

[50] N.X. Landén, D. Li, M. Ståhle, Transition from inflammation to proliferation: a critical step during wound healing, Cellular and Molecular Life Sciences 73 (2016) 3861–3885.

[51] M.G. Jeschke, M.E. van Baar, M.A. Choudhry, K.K. Chung, N.S. Gibran, S. Logsetty, Burn injury, Nature reviews Disease primers 6(1) (2020) 11.

[52] S. Werner, T. Krieg, H. Smola, Keratinocyte–fibroblast interactions in wound healing, Journal of investigative dermatology 127(5) (2007) 998–1008.

[53] M.V. Plikus, X. Wang, S. Sinha, E. Forte, S.M. Thompson, E.L. Herzog, R.R. Driskell, N. Rosenthal, J. Biernaskie, V. Horsley, Fibroblasts: Origins, definitions, and functions in health and disease, Cell 184(15) (2021) 3852–3872.

[54] B. Russo, N.C. Brembilla, C. Chizzolini, Interplay between keratinocytes and fibroblasts: a systematic review providing a new angle for understanding skin fibrotic disorders, Frontiers in immunology 11 (2020) 648.

[55] Z. Deng, T. Fan, C. Xiao, H. Tian, Y. Zheng, C. Li, J. He, TGF-β signaling in health, disease and therapeutics, Signal transduction and targeted therapy 9(1) (2024) 61.

[56] K.A. Cook, E. Martinez-Lozano, R. Sheridan, E.K. Rodriguez, A. Nazarian, M.W. Grinstaff, Hydrogels for the management of second-degree burns: currently available options and future promise, Burns & trauma 10 (2022) tkac047.

[57] N. Mohamad, E.Y.X. Loh, M.B. Fauzi, M.H. Ng, M.C.I. Mohd Amin, In vivo evaluation of bacterial cellulose/acrylic acid wound dressing hydrogel containing keratinocytes and fibroblasts for burn wounds, Drug delivery and translational research 9 (2019) 444–452.

[58] C.M. Abreu, R.P. Pirraco, R.L. Reis, M.T. Cerqueira, A.P. Marques, Interfollicular epidermal stem-like cells for the recreation of the hair follicle epithelial compartment, Stem cell research & therapy 12 (2021) 1–12.

[59] I. Hwang, K.-A. Choi, H.-S. Park, H. Jeong, J.-O. Kim, K.-C. Seol, H.-J. Kwon, I.-H. Park, S. Hong, Neural stem cells restore hair growth through activation of the hair follicle niche, Cell Transplantation 25(8) (2016) 1439–1451.

[60] R.T. Kendall, C.A. Feghali-Bostwick, Fibroblasts in fibrosis: novel roles and mediators, Frontiers in pharmacology 5 (2014) 123.

[61] J. Roman, Fibroblasts—warriors at the intersection of wound healing and disrepair, Biomolecules 13(6) (2023) 945.

[62] A.D. Widgerow, S.G. Fabi, R.F. Palestine, A. Rivkin, A. Ortiz, V.W. Bucay, A. Chiu, L. Naga, J. Emer, P.E. Chasan, Extracellular Matrix Modulation: Optimizing Skin Care and Rejuvenation Procedures, Journal of drugs in dermatology: JDD 15(4 Suppl) (2016) s63–71.

[63] K.W. Finnson, S. McLean, G.M. Di Guglielmo, A. Philip, Dynamics of transforming growth factor beta signaling in wound healing and scarring, Advances in wound care 2(5) (2013) 195–214.

