## Supplementary material for "Topical Delivery of 4-Aminopyridine Enhances Skin Regeneration in Burn Wounds": Supplimental File

<u>**Supplementary Data**</u>

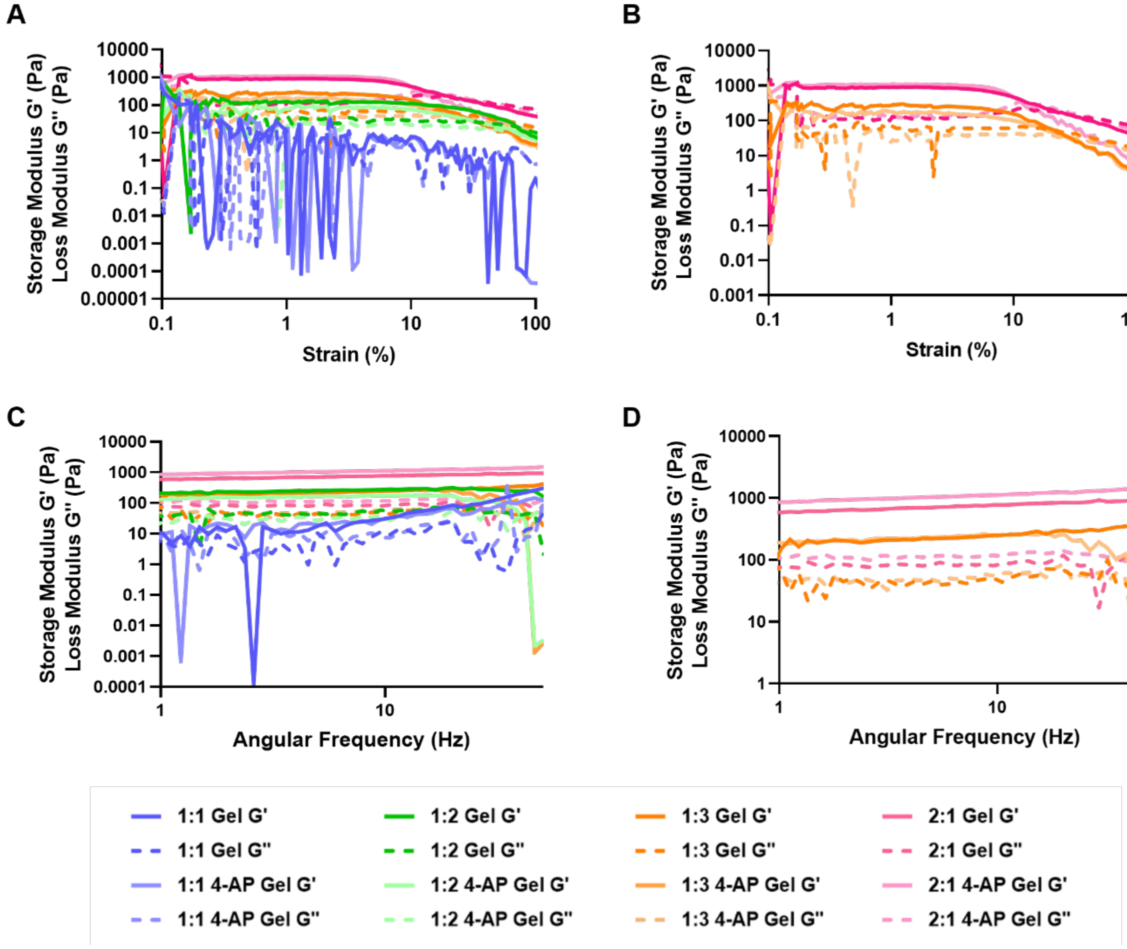


**Fig. S1.** Rheological characterization of formulated gels performed at room temperature (A and B) Amplitude sweep. (C and D). Frequency sweep.

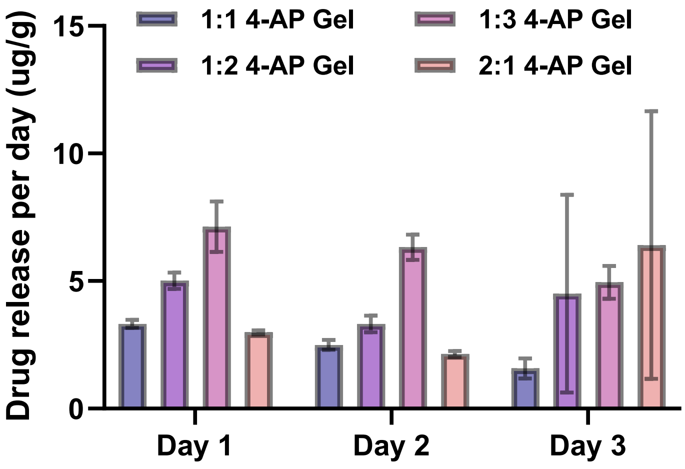


**Fig. S2.** 4-Aminopyridine drug release from different composition of gels

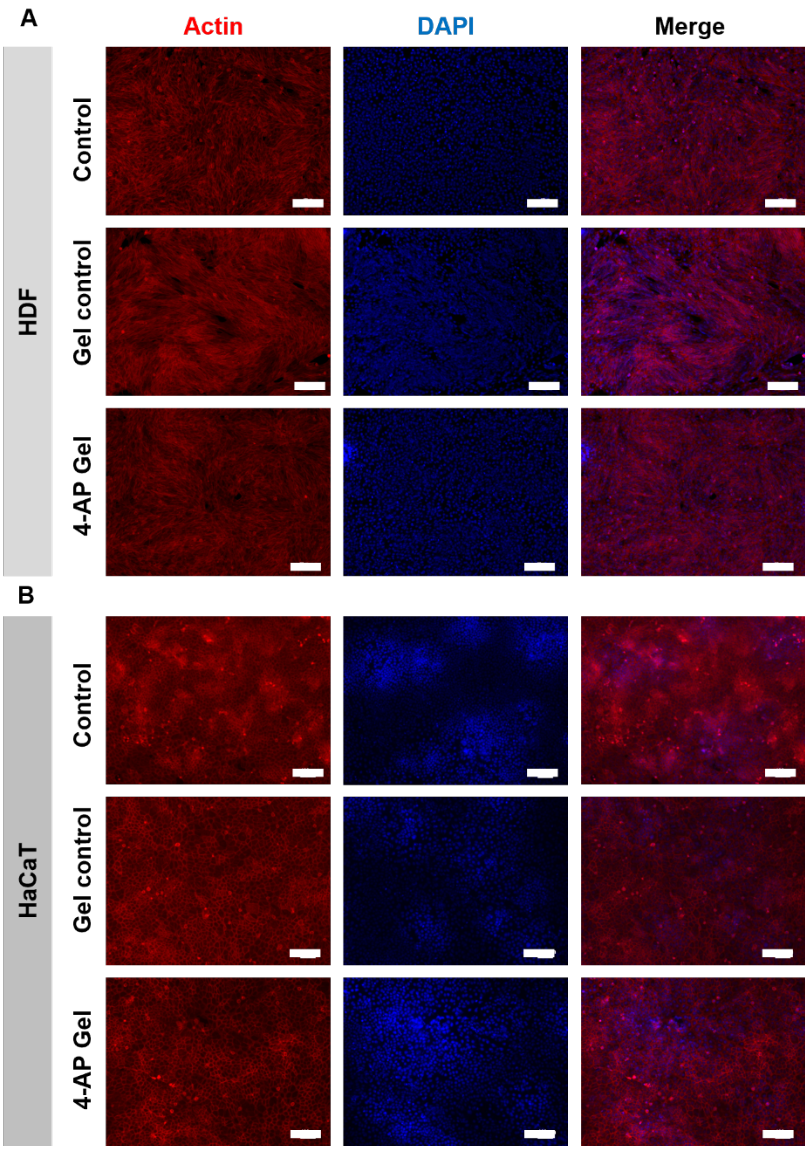


**Fig. S3.** Actin cytoskeleton staining after 3 days of culture with control and 4-AP gels, red fluorescence depicts cytoskeleton and blue represents nuclei of the cell. (A) Actin staining of HDF. (B) Actin staining of HaCaT. Scale bar 200 μm.

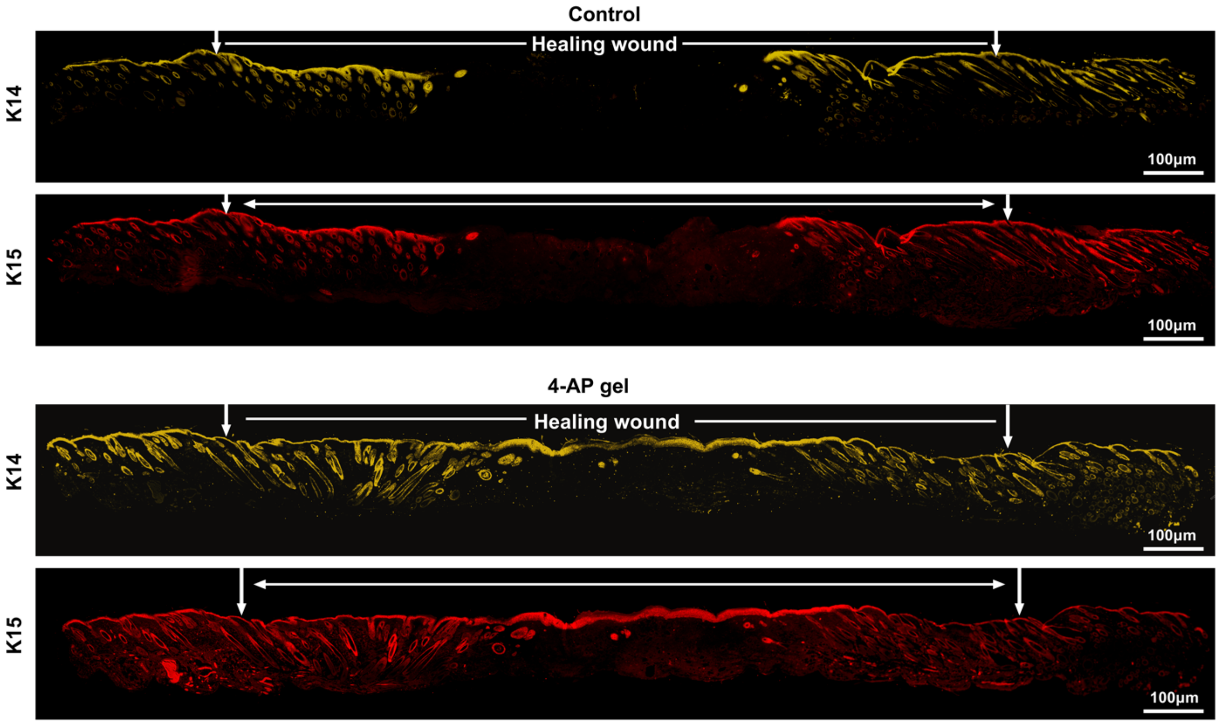


**Fig. S4.** Representative tile scanning IF images of K14 and K15.

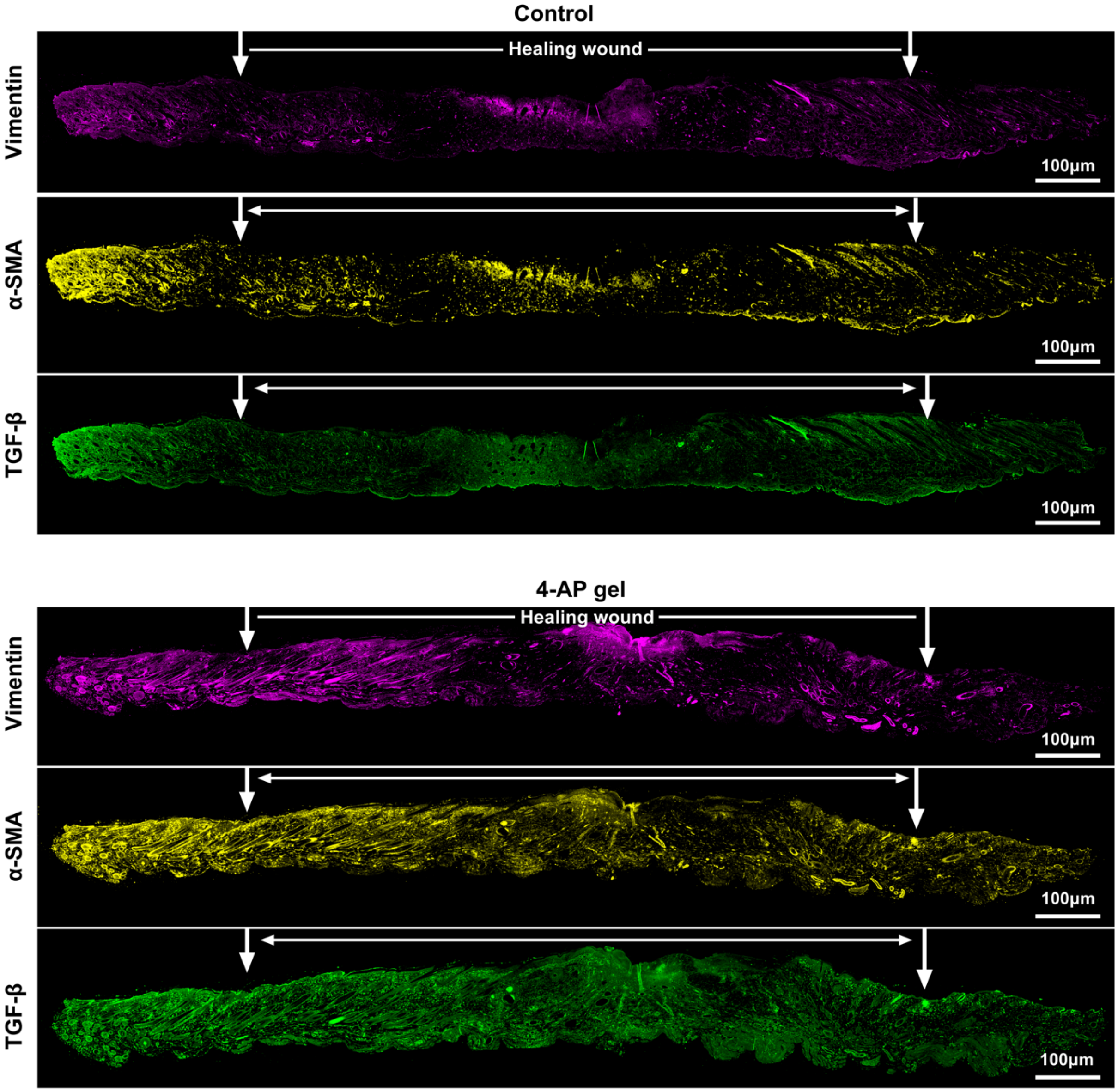


**Fig. S5.** Representative tile scanning IF images of vimentin, α-SMA, and TGF-β.

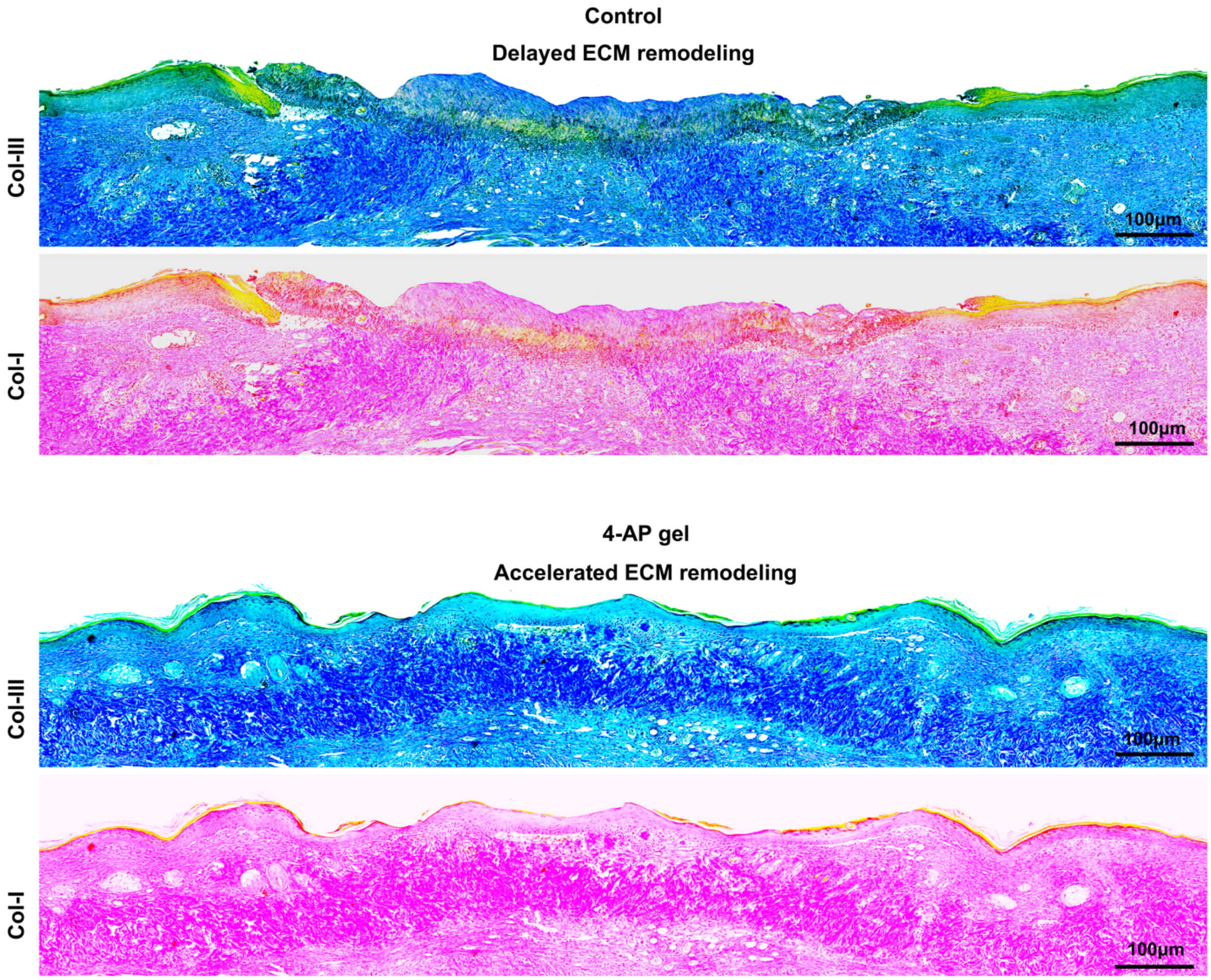


**Fig. S6.** Representative split channel images of collagen types I (pink to red) and III (blue) obtained from the ImageJ.

**Table S1.** Primers used for gene expression analysis

| **Gene** | **Primer sequence (5ʹ-3ʹ)** | |
|---|---|---|
| | **Forward** | **Reverse** |
| **K14** | GTTTTCGGCCTCTCGCTAGTT | CTGTTCCCGGTCTTGAACC |
| **K15** | AACTCTGGAAACTACTCCTCACA | AGGGCAAGTGGCCTTCAAG |
| **Vimentin** | CGGCTGCGAGAGAAATTGC | CCACTTTCCGTTCAAGGTCAAG |
| **α-SMA** | GACAGGGCAATCACCGTCTTC | CGAGAGCGCAGATTTTCCTCA |
| **TGF-β** | AGACCACATCAGCATTGAGTG | GGTGGCAACGAATGTAGCTGT |
| **Col-I** | GCTCCTCTTAGGGGCCACT | CCACGTCTCACCATTGGGG |

| | | |
|---|---|---|
| **Col-III** | AGATGAGGCGGAAACTCAAGT | AGCCCTTTAGGAAGAGGTGTT |
| **GAPDH** | TGGATTTGGACGCATTGGTC | TTTGCACTGGTACGTGTTGAT |
